# Cynipid wasps systematically reprogram host metabolism and restructure cell walls in developing galls

**DOI:** 10.1101/2023.05.22.540048

**Authors:** Kasey Markel, Vlastimil Novak, Benjamin P. Bowen, Yang Tian, Yi-Chun Chen, Sasilada Sirirungruang, Andy Zhou, Katherine B. Louie, Trent R. Northen, Aymerick Eudes, Henrik V. Scheller, Patrick M. Shih

## Abstract

Many insects have evolved the ability to manipulate plant growth to generate extraordinary structures called galls in which insect larva can develop while being sheltered within and feeding on the plant. In particular, Cynipid (Hymenoptera: Cynipidae) wasps have evolved to form some of the most morphologically complex galls known and generate an astonishing array of gall shapes, colors, and sizes. However, the biochemical basis underlying these remarkable cellular and developmental transformations remains poorly understood. A key determinant in plant cellular development is the deposition of the cell wall to dictate the physical form and physiological function of newly developing cells, tissues, and organs. However, it is unclear to what degree cell walls are restructured to initiate and support the formation of new gall tissue. Here, we characterize the molecular alterations underlying gall development using a combination of metabolomic, histological, and biochemical techniques to elucidate how leaf cells are reprogrammed to form galls. Strikingly, gall development involves an exceptionally coordinated spatial deposition of lignin and xylan to form *de novo* gall vasculature. Our results highlight how Cynipid wasps can radically change the metabolite profile and restructure the cell wall to enable the formation of galls, providing new insights into the mechanism of gall induction and the extent to which plants can be entirely reprogrammed to form novel structures and organs.

## Introduction

Diverse organisms from fungi and bacteria to plants and insects have independently evolved the ability to manipulate the growth of plants to their advantage, forming abnormal structures referred to as galls. Galls induced by bacteria and fungi are generally morphologically simple and often referred to simply as ‘tumors’, whereas galls induced by insects are sometimes intricately and precisely structured (1) and have fascinated naturalists since the time of ancient Greece (2, 3).

While the exact mechanisms of gall induction remain largely unknown, chemical signals from the insect causing plant growth reprogramming has been the primary working hypothesis since the time of Charles Darwin, who presents “the poison secreted by the gall-fly produces monstrous growths on the wild rose or oak-tree” as one of the final arguments suggesting all plants (and in fact all life) share common ancestry (4). Interestingly, some gall inducers create galls on several species of host plants, and in these cases the gall morphology is remarkably similar (5). This demonstrates that the gall is an extended phenotype of the gall inducer (6, 7), which exerts greater control over gall morphology than the plant host. The genes underlying this extended phenotype in insect gall inducers remain almost entirely unknown, but the phenotype itself is striking: precise control over host growth, metabolism, and structure.

Deciphering of the mechanisms of gall induction has been a longstanding goal of gall research (1, 8). One major theory is that gall inducers synthesize plant hormones or hormone analogs, the local concentration gradients of which play a key role in gall development (9, 10). Synthesis of plant hormones – principally auxin and cytokinin – is known to be a key component in the generation of the simple galls induced by *Agrobacterium* (11). Nonetheless, hormones likely play some role in insect gall induction, a hypothesis supported by the detection of high concentrations of plant hormone analogues in gall tissue (12), though studies of other gall types have found galls to be auxin-depleted compared to normal tissue (13). Even more conspicuous evidence comes from the ability of some gall-inducing insects to synthesize plant hormones such as auxin and most likely cytokinin (13–15). Taken together, the balance of evidence suggests that phytohormone synthesis plays a role in the induction of at least some galls, but the exact mechanism is unclear and there are almost certainly other unknown elements to the induction of galls. In particular, simple concentration gradients of hormones are insufficient to explain the morphological diversity and complexity of insect-induced galls; thus there is a need to discover, study, and expand our understanding of the many non-phytohormone compounds that may contribute to the development and morphology of complex galls.

Because cell walls physically surround and constrain plant cells, any new growth such as the development of a gall requires breakdown, remodeling, and/or new deposition of cell wall material. As such, cell wall remodeling is key to organogenesis, such as the generation of new leaves (16). Despite the central role cell walls play in determining the structure and function of plant tissues, little is known about how plant cell walls are modified during the development of insect galls. Previous qualitative studies have shown changes in lignin (17), tannins (18)), and several polysaccharides (19, 20); however, there is a need for a more global understanding of all the metabolic changes underlying the transformation of the plant cell metabolites and cell walls during the formation of galls. Similarly, a more detailed spatial understanding of the molecular alteration in plant cell walls associated with gall induction may reveal unique insights into the relatively unexplored interplay between host cell reprogramming and cell wall remodeling.

Among the most morphologically complex and charismatic galls are those induced by *Cynipid* wasps on oak trees (21). Over 1300 species of cynipid wasps have been described (22), and many species alternate between a sexual and parthenogenic generation, each of which produces a distinct gall type (23). The diversity and morphological complexity of cynipid galls make them an excellent system to study the morphological, metabolic, and cell wall changes associated with gall induction. Recent molecular biology research on cynipid galls has been largely limited to transcriptomics studies (24, 25). These analyses have shed some light on the question of how cynipid wasps hijack the gene expression machinery of plants, but the metabolic consequences of these changes in gene expression remain largely unexplored. However, because insect gall induction is believed to be dependent on the generation of gradients of signaling molecules such as phytohormones and requires changes to cell wall structure and composition without obvious mRNA correlates, transcriptomics alone cannot tell the whole story. To provide a more comprehensive understanding of oak gall development, we perform a detailed analysis of the biochemical changes associated with gall induction and the concurrent alterations to cell wall structure and composition.

## Results

### Morphological characterization of two distinct gall types

We collected cone galls induced by *Andricus kingi* and urchin galls induced by *Antron douglasii* from the leaves of the valley oak *Quercus lobata* in and near the UC Davis arboretum (trees sampled shown in Supplemental Figure 1, sampling dates and other galls identified shown in Dataset S1). Cone galls (Figure 1A) are cone shaped, usually red but often white along one side, and approximately 5 mm across at maturity. Urchin galls (Figure 1B) are rarer and larger, light purple in color, and urchin-shaped with 5-15 spikes. Both are attached to the leaf by a thin (<200 µm) section of tissue that projects orthogonal to the plane of the leaf blade and defines the axis of rotational symmetry for cone galls and approximate symmetry for the urchin galls.

**Figure 1:**
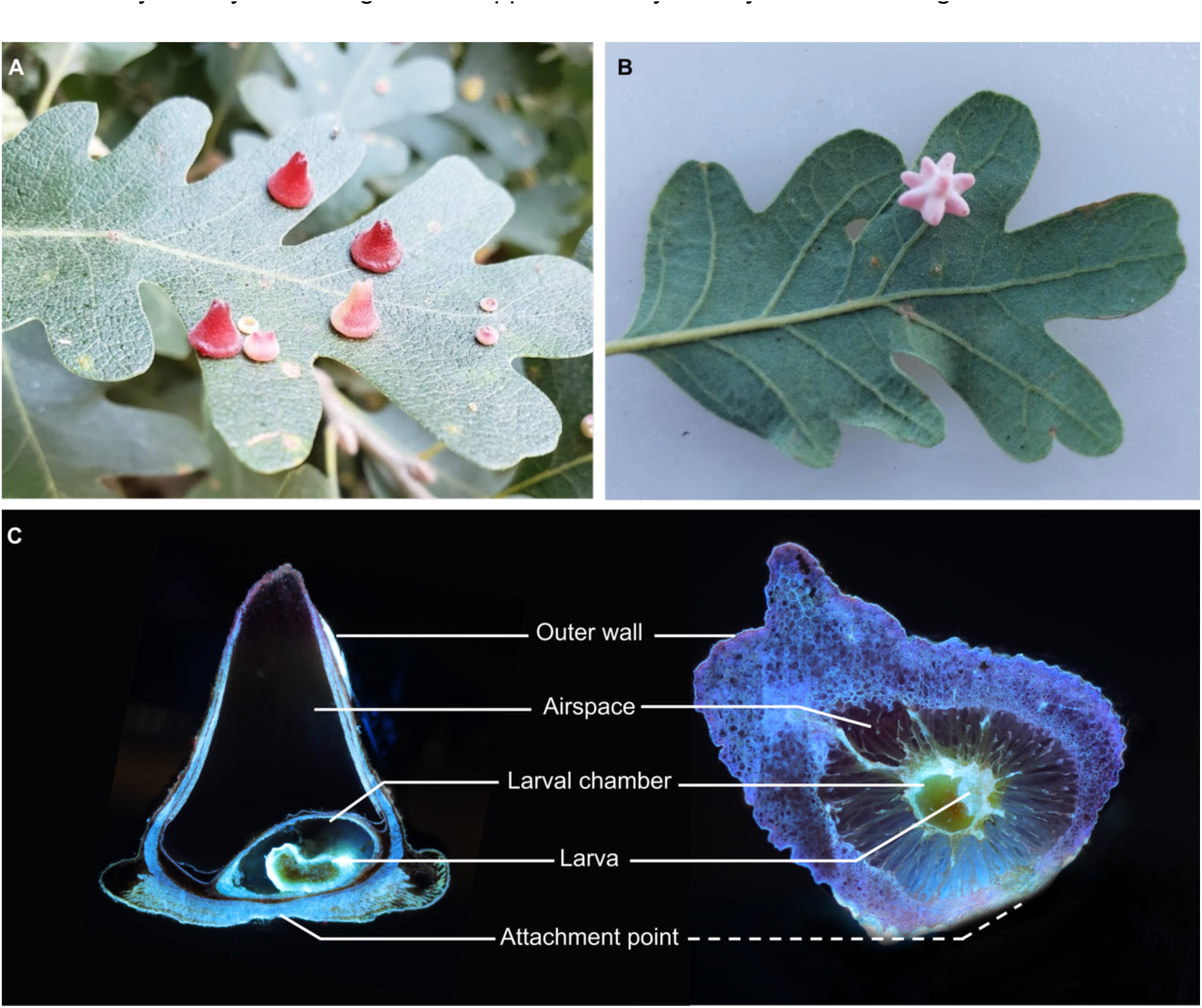
Comparison of anatomical features shared across morphologically disparate galls. A: Cone galls induced by *Andricus kingi*. B: Urchin gall induced by *Antron douglasii*. C: Longitudinal sections of cone (left) and urchin (right) galls, imaged with laser ablation tomography. Dashed line to urchin gall attachment point indicates attachment point is out of plane and location is approximate.

Galls induced by cynipid wasps are complex three-dimensional structures; however, the vast majority of studies have been constrained to two-dimensional sections, which has been insufficient to comprehensively characterize the relationship between plant and insect tissue. To fill this gap, we used laser ablation tomography (LAT) to generate high-resolution three-dimensional reconstructions of galls consisting of thousands of two-dimensional slices with ∼8 μm resolution (Figure 1C, Supplemental Figure 2). Three-dimensional models reveal the internal structure of cone (Supplemental Videos 1 and 2) and urchin (Supplemental Video 3) galls. The *Andricus kingi* larva within the cone galls is highly autofluorescent, facilitating easy discrimination between larval and plant tissue. Surprisingly, we found the larva in different orientations in the two cone galls imaged, with the long axis aligned to the longitudinal axis of the cone gall in one case and orthogonal in the other. This variation in the orientation of the larval chamber in conjunction with the tight conservation of the overall gall structure suggests something other than the larva provides the “orientation lodestar”, most likely the attachment point to the leaf.

While the morphology of both types of gall is very different, they both consist of a relatively thick outer wall of plant cells, an airspace, and an inner layer of plant cells surrounding a fluid-filled cavity which houses the developing larva. The urchin gall larval chamber is suspended by thin strands of plant tissue in the center of the airspace, providing thermal insulation. The thick exterior wall and airspace have been demonstrated to be important for protection of the larva from the elements (26) and hypothesized to be important for protection from predators and parasitoids (21). An evolutionary arms race between gall inducers and these natural enemies is likely a contributing source of the tremendous variation in cynipid gall morphology.

The three-dimensional models show that plant epidermal cells surrounding the larval chamber are patterned in a smooth ovate structure. While the galls imaged with LAT were relatively mature, the insect larva remained fairly undeveloped, and likely incapable of chewing through the plant cells surrounding the chamber. However, they were within an order of magnitude of the size of the adult wasps, which means they had almost certainly grown quite substantially to reach their current size. These facts together support the hypothesis that up to and including this gall developmental stage, insect larvae are absorbing nutrients through the fluid within the larval chamber, much like other animals feed on energy reserves within an egg or plant seedlings feed on endosperm, and in contrast to the mechanical chewing found in galling thrips (27) and during the final stages of cynipid development (28). The fluid of the larval chamber is most likely to be translocated photosynthate and nutrients, highlighting the importance of vasculature to support proper gall development.

### Metabolomic profiles of different gall types are distinct and unique

We examined the metabolomic profiles of the two gall types, looking for common patterns that may suggest the homologously shared mechanism of gall induction as well as differences that may explain the differences in gall morphology. Previous research has focused either on a small number of pre-selected metabolites (29–31) or on the transcriptional profile of galls (24, 25). We utilized untargeted metabolomics to identify metabolite differences between gall and normal leaf tissues, enabling detection of thousands of mass features between samples. The most recent common ancestor of the two species of gall wasp studied most likely also induced galls (32), and therefore shared changes in the metabolomic profile of the two galls may suggest key elements of the basic mechanism of gall induction, whereas differences between the two galls may be either a cause or result of more idiosyncratic elements of gall structure or random changes due to drift.

Metabolic changes associated with initial induction of galls are expected to be especially pronounced during the early stages of gall development. Therefore, cone and urchin galls were subdivided into 5 and 4 developmental stages respectively using mass as a proxy for developmental stage. To our knowledge, the resulting dataset is the first metabolomic analysis of cynipid gall development incorporating a developmental time-series design. We obtained 8690 mass features; the full datasets for positive and negative mass spectrometry modes are available as Dataset S2 and S3, respectively, heat map of mass feature peak height available in Supplemental Figure 3. Principal component analysis demonstrated mass feature composition was distinct for leaf, urchin gall, and cone gall samples (Figure 2A). The majority (63%) of these mass features were shared between at least two sample types, and 29% were shared among all three (Figure 2B).

**Figure 2:**
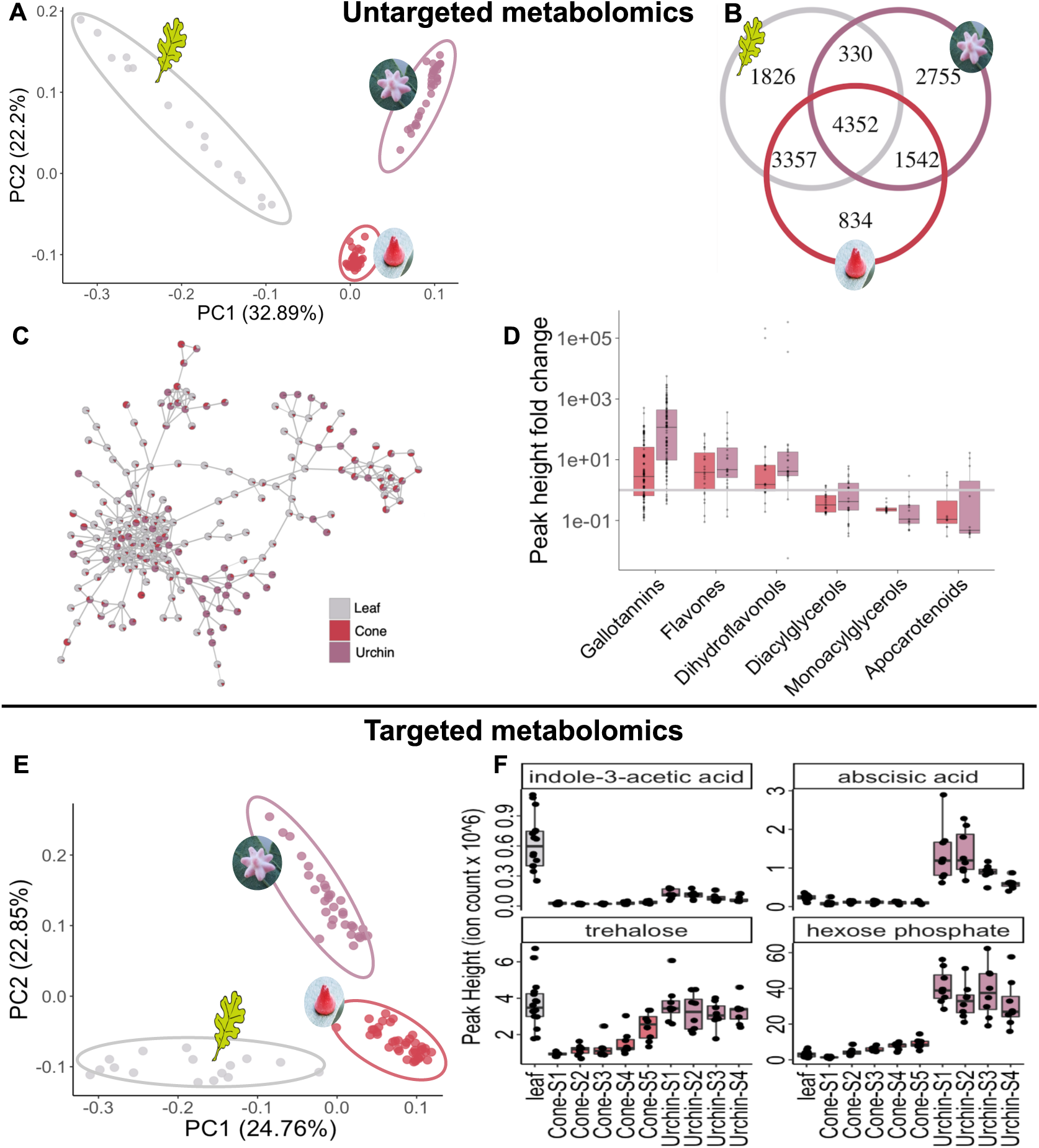
Galls are metabolically distinct from each other and leaf tissue. A: Principal component analysis of all 8690 mass features recorded in positive mode in untargeted metabolomics B: Venn Diagram of the mass features present in each sample type in untargeted metabolomics. C: Molecular networking. Each node is a mass feature, each edge indicates a cosine score of fragmentation pattern of at least 0.7, nodes are pie charts indicating relative peak heights between the three sample types. D: Natural products classes of putative identifications of mass features in untargeted metabolomics. E: Principal component analysis of all 209 metabolites positively identified with mass-charge ratio, secondary fragmentation pattern, and retention time confirmed against a library for the same instrument in positive and negative mode, removing whichever was lower to generate a nonredundant dataset. F: Metabolite data for leaf and several growth stages of each type of gall for two hormones and two sugars. MS-MS mirror plots with more precise identification information abscisic acid, trehalose, and hexose phosphate are available in Supplemental Figures 6, 7, and 8 respectively. Indole-3-acetic acid peak height was too low to trigger MS-MS, identification was based on retention time, m/z ratio, and other mass feature characteristics shown in Dataset S4.

We performed network analysis using GNPS, which revealed that mass features overrepresented in particular sample types often clustered, demonstrating similar classes of compounds were enriched in specific galls (Figure 2C, Supplemental Figure 3). Several interactive networks are available online at NDExbio – further described in methods. We used m/z ratio and networking to generate putative identifications for each mass feature and used NPClassifier to categorize them (33), revealing increases in several expected compound classes such as gallotannins (whose name derives from ‘gall’) in gall tissue compared to leaf. We also observed an increase in two flavonoid categories and a decrease in two acylglycerol categories as well as apocarotenoids (Figure 2D, Supplemental Figure 5). Our finding of increased flavonoid accumulation corroborates a recent report of upregulation of flavonoid biosynthetic genes in cynipid-oak galls (25), which also may be the underlying basis of the pigmentation of the galls themselves. The decrease in acylglycerols is consistent with the same study observing that 2 of the top 50 upregulated genes were annotated as “hydrolysis of fatty acids.” It has been proposed that fatty acids are converted into sugars to feed the growing larva (25). Finally, cynipid galls have previously been shown to contain lower concentrations of chlorophyll and carotenoids (34, 35), suggesting reduced photosynthesis as an explanation for the reduction in apocarotenoids observed here.

To provide a more detailed and quantitative understanding of metabolite changes, we next performed targeted metabolomics based on a library of standards with known retention time and fragmentation data to identify specific metabolites that broadly cover a wide sampling of primary metabolism and many core plant metabolites. Targeted metabolomic analyses combined the positive and negative mode datasets by choosing whichever had higher peak height (following methodology from (36)), resulting in a non-redundant dataset of 209 metabolites with confidence score of at least “Level 1”, meaning at least two independent and orthogonal data are used to confirm metabolite identity (37). Identification evidence including MS1, MS2, and chromatographic peak comparisons are available as Dataset S4 and S5 for positive and negative modes respectively. Principal component analysis of this stringently curated dataset revealed a distinct separation of sample types (Figure 2E), the full dataset is available as Dataset S6. Additional principal component analyses of the growth stages of each type of gall reinforce clear distinction between gall and leaf metabolites and show some clustering by gall growth stage (Supplemental Figure 3).

Of these 209 identified metabolites from targeted metabolomics, 43 had peak heights in urchin galls greater than four times higher than the leaf average, and 22 had peak heights in cone galls greater than four times higher than the leaf average. Of these highly gall-abundant metabolites, 11 were enriched in both gall types, much more than would be predicted if peak height were independent in both gall types (p = 0.001, hypergeometric test). Peak height data for the 54 metabolites >4x higher peak height in galls compared to leaf tissue is available in Dataset S7, these metabolites are candidates for either causes or conserved metabolic effects of the gall induction process, and may be useful leads for future efforts to determine the mechanism of gall induction. We used NPClassifier to classify all 209 metabolites by pathway and evaluated whether any pathways were overrepresented among the metabolites enriched in galls. For both gall types, there were fewer fatty acids than chance (p=0.052 for cone galls, p = 0.024 for urchin galls, hypergeometric test), supporting the results from the untargeted metabolomics.

### Conserved metabolite changes across different galls reveals drastic changes in plant hormone and sugar concentrations

We next examined the concentration of plant hormones detected in the metabolomic analyses, which have been hypothesized to play important roles in the gall induction process. Structurally complex galls can be thought of as a novel organ functioning for the benefit of the gall inducer, and hormone concentration gradients are known to be central to the growth of organs such as leaves (38), flowers (39, 40), and fruits (39). Interestingly, the transcriptomic profile of galls induced by phylloxera on grape leaves shares many similarities to the transcriptome of fruits (41).

We found major differences in the concentration of auxin (indole-3-acetic acid) and abscisic acid between galls and normal leaf tissue (Figure 2F). Existing literature shows that auxin and cytokinin are sometimes increased and sometimes decreased in gall tissue compared to normal plant tissue, suggesting there may be multiple separate mechanisms of plant growth manipulation used by different groups of gall inducers (reviewed in (42)). This is perhaps not surprising given that the gall-inducing habit has evolved independently many times in separate lineages (43). In both cone and urchin galls, we see a massive decrease in the concentration of auxin (Figure 2F). This is somewhat surprising given the relatively low baseline levels of auxin in the middle of a leaf lamina (44), and even more surprising in light of the fact that RNAseq of a closely related cynipid-induced oak gall showed upregulation of auxin-response genes (24). While it is possible that these discordant results reflect different ground truths in these closely-related cynipid galls, it is also possible that upregulation of auxin biosynthetic genes does not result in increased auxin accumulation, highlighting a potential pitfall of interpretations of small molecule concentration solely made by transcript levels without direct biochemical measurement.

A recent gall tissue-specific RNAseq study found upregulation of auxin biosynthetic genes only in the larval chamber tissue, which comprises a relatively small portion of the total gall biomass, with low expression of auxin responsive genes throughout the remainder of the gall (25). This finding may help reconcile the seemingly contradictory results: while auxin is involved somehow in the gall induction process, if only the gall larval chamber contains high concentrations of auxin, then depending on the mass ratio of the larval chamber compared to the exterior of the gall we would expect to see some reports of higher auxin concentration and some reports of lower concentration within galls, which is indeed what has been reported (42).

Abscisic acid concentration is increased in urchin galls, but not cone galls (Figure 2F). Abscisic acid is often associated with stress, and has been shown to increase in response to attempted gall induction on resistant plants, while remaining constant between gall and normal tissue in susceptible plants (45). In another gall system, abscisic acid was reported to be decreased in gall tissue compared to normal plant tissue (46). In light of these diverging results among very phylogenetically distant gall systems, it is interesting to see different behavior in abscisic acid response even among two closely related galls on the same plant host. It is also worth noting that the only mass features identified as apocarotenoids increased in gall tissue in Figure 2D were putatively identified as abscisic acid. Since this was an independant mass spectrometry run, that both strengthens the results from this targeted analysis and their removal from the apocarotenoid class (of which abscisic acid is clearly a non-central example) strengthens the finding that apocarotenoids are depleted in gall tissue.

We also observed a striking pattern in the concentration of trehalose, a disaccharide known to play important signaling and regulatory roles. Trehalose plays a stress signaling role in plants, and exogenous application of trehalose induces resistance against pathogens (47). The massive reduction of trehalose concentration in cone galls may suggest the wasps are silencing this defense response. Trehalose also plays important roles in insects; it is a major circulating carbohydrate in the hemolymph (48), as well as a regulator of long-term hibernation-like states (49). Therefore, further research is necessary to fully understand the significance of the trehalose reduction in gall tissue.

We next examined hexose phosphates, central metabolic intermediates which are a primary output of photosynthesis and primary input into cell wall assembly. Hexose phosphates are substantially enriched in all surveyed developmental stages of urchin gall tissue compared to leaf, but remain constant at leaf-like levels in cone galls (Figure 2F). On average, hexose phosphate levels in urchin galls are over ten times higher than the leaf baseline. In general, the majority of hexose phosphates are destined for generation of starch or cell wall polysaccharides, suggesting the rerouting of metabolism to support gall development and larval feeding.

### Gall cell layers are chemically distinct and highly lignified suggestive of de novo vascularization

Though the three-dimensional models generated by laser ablation tomography offer unique structural insights, they lack the chemical information. Metabolomic analysis offers chemical information, but without spatial data. To address the intersection of these interests, we turned to histochemical staining. Histochemical staining is a standard approach to identifying plant tissue types, yet there are no published micrographs of either of the galls studied here. Therefore, we next used a series of classic plant histology stains on cone galls (chosen for microscopy as they were more abundant) to examine the chemical composition and distribution to better understand the chemical changes associated with gall development. Safranin O, congo red, Mäule stain, cellulose azure, orange G, FastGreen FCF, and aniline blue failed to show any interesting spatial patterns within the gall material (Supplemental Figure 9). Toluidine blue was useful for generating contrast to determine cell wall morphology and differentiate cell layers (Supplemental Figure 10). Weisner reagent (phloroglucinol + HCl) revealed the most striking spatial pattern, demonstrating tight spatial regulation of lignin deposition in gall tissue (Figure 3A, B). Two sclerenchyma cell layers are strongly stained (Supplemental Figure 10A), and the central sponge layer between them contains bundles of 4-9 cells in cross section with moderate lignification, which is suggestive of vasculature.

**Figure 3:**
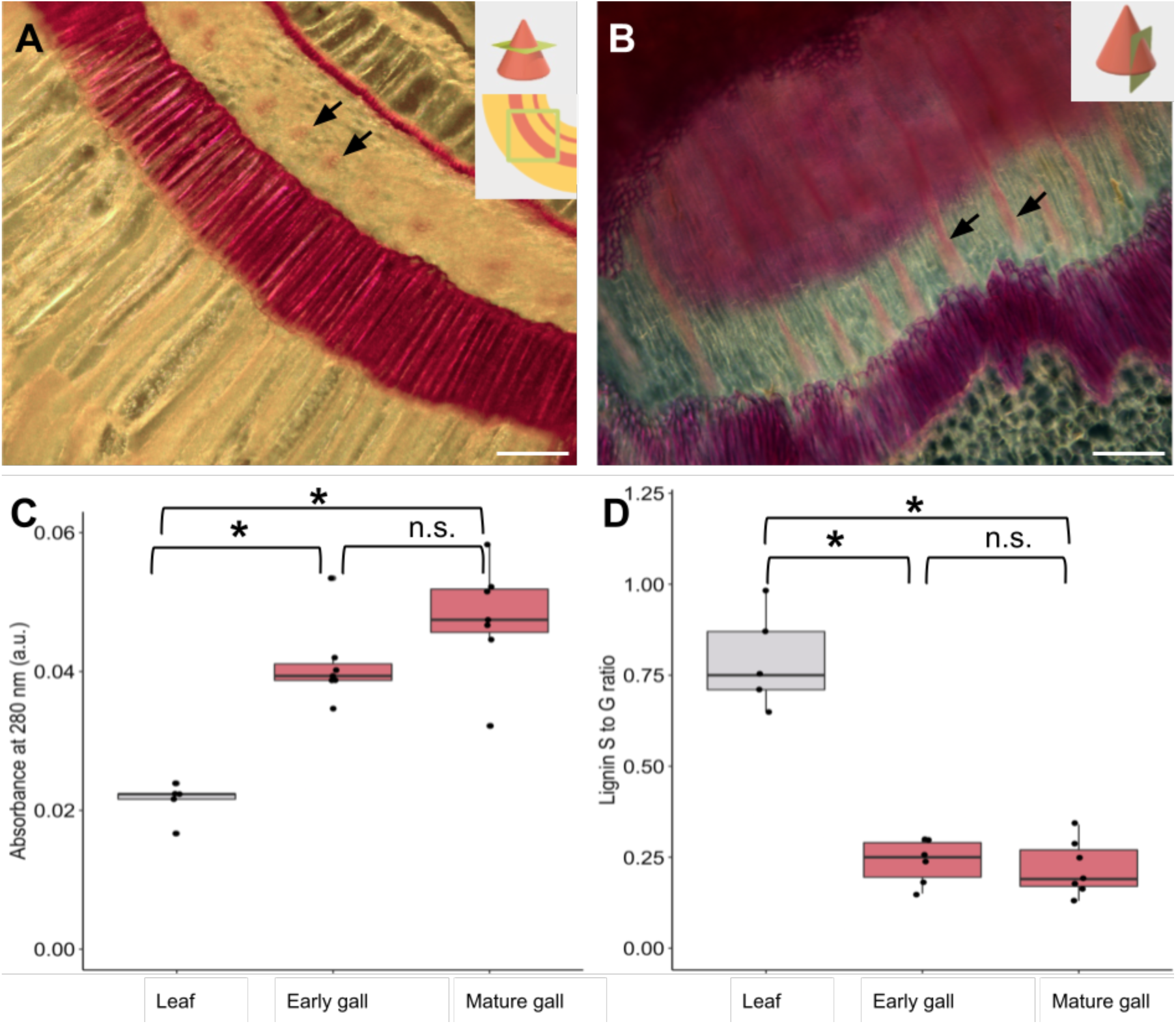
Lignin deposition in cone galls is spatially coordinated in a gall-specific pattern. A: Transverse section of cone gall stained with Weisner stain, showing two heavily lignified cell layers and one cell layer containing bundles of 4-9 highly lignified cells (arrows). Scale bar = 100 µm. B: Darkfield image of tangential longitudinal section of gall stained with Weisner stain, which stains heavily lignified tissue pink. The same two heavily lignified cell layers are visible, as well as the small moderately lignified bundles (arrows), now in longitudinal section. Scale bar = 100 µm. C: Lignin concentration in leaf tissue, early-development cone galls, and mature cone galls, as determined by thioglycolic acid assay. Asterisks indicate p < 0.05 by Kruskal-Wallis test with Benjamini-Hochberg correction for multiple comparison, n.s. Indicates p > 0.05. D: Lignin subunit S to G ratio as determined by pyro-GC MS. Asterisks indicate p < 0.05 by Kruskal-Wallis test with Benjamini-Hochberg correction for multiple comparison, n.s. Indicates p > 0.05

Weisner staining revealed large amounts of lignin, but histological studies cannot provide an accurate quantification of these chemical changes. To fill this gap, we used the thioglycolic acid assay to quantify lignin in leaf and gall tissue, comparing leaf tissue against young or mature cone galls, as shown in Figure 3C. Cone galls are substantially more lignified than leaf tissue (p=0.0025, Kruskal-Wallis test with Benjamini-Hochberg correction for multiple comparisons), but the two developmental stages are statistically indistinguishable from each other. In light of this substantial increase in lignin levels, we asked whether the lignin monomeric composition was altered as well using pyrolysis gas chromatography coupled to mass spectrometry (pyro-GC MS). Lignin polymers are composed of three subunits, namely syringyl (S), guaiacyl (G), and *p*-hydroxyphenyl (H), which polymerize with a complex branched structure that is highly resistant to degradation (50, 51). Lignin associated with fiber cells tends to contain a higher fraction of S subunits, whereas vascular elements contain more G subunits (52). The S to G ratio was substantially lower in both stages of gall tissue (p = 0.011 and 0.0076 for early and mature galls respectively, Kruskal-Wallis test with Benjamini-Hochberg correction for multiple comparisons, full data for all lignin-derived fragments available in Dataset S8) compared to leaf tissue, which also supports generation of vasculature in the galls.

To our knowledge, this is the first description of *de novo* generation of vascular elements in insect galls, and our findings are in contrast to a detailed analysis of another cynipid-induced gall, where *de novo* production of vasculature was specifically ruled out (53), suggesting neovascularization only occurs in some types of cynipid galls. The importance of obtaining access to the plant vascular system has long been recognized as important for the growth and success of galling insects (54), but previous studies have shown modifications of existing vasculature rather than *de novo* vascularization (53, 55). In contrast, the histological evidence here demonstrates gall generation involves coordinated, spatially organized generation of *de novo* vasculature.

### Cell wall remodeling is associated with gall formation

The cell wall plays an integral role in defining the form and function of plant cells. Although we had already observed changes in lignin composition and deposition, the majority of the cell wall is composed of polysaccharides (56). Changes in polysaccharide content can drastically alter the biochemical, physical, and ultimately physiological role of plant cells and tissues. A large portion of plant sugars are ultimately sequestered in the cell wall as polysaccharides, and in conjunction with lignin comprise the primary physical support structure of plant organs. Since metabolomics revealed differences in hexose phosphate concentrations, we reasoned that this could lead to changes in the monosaccharide composition of the cell walls. Indeed, cell wall composition varied wildly between galls and the leaf tissue from which they arise, as shown in Figure 4A. Notably, xylose residues were extremely abundant in gall tissue, to the extent that all other monosaccharide signals are largely suppressed, and surprisingly, xylose accounts for over 75 percent of all hydrolyzed cell wall monosaccharides in cone gall samples. It should be noted that the cell wall polysaccharide hydrolysis method employed leaves cellulose intact and measures the monosaccharide composition of all non-cellulosic cell wall polysaccharides.

**Figure 4:**
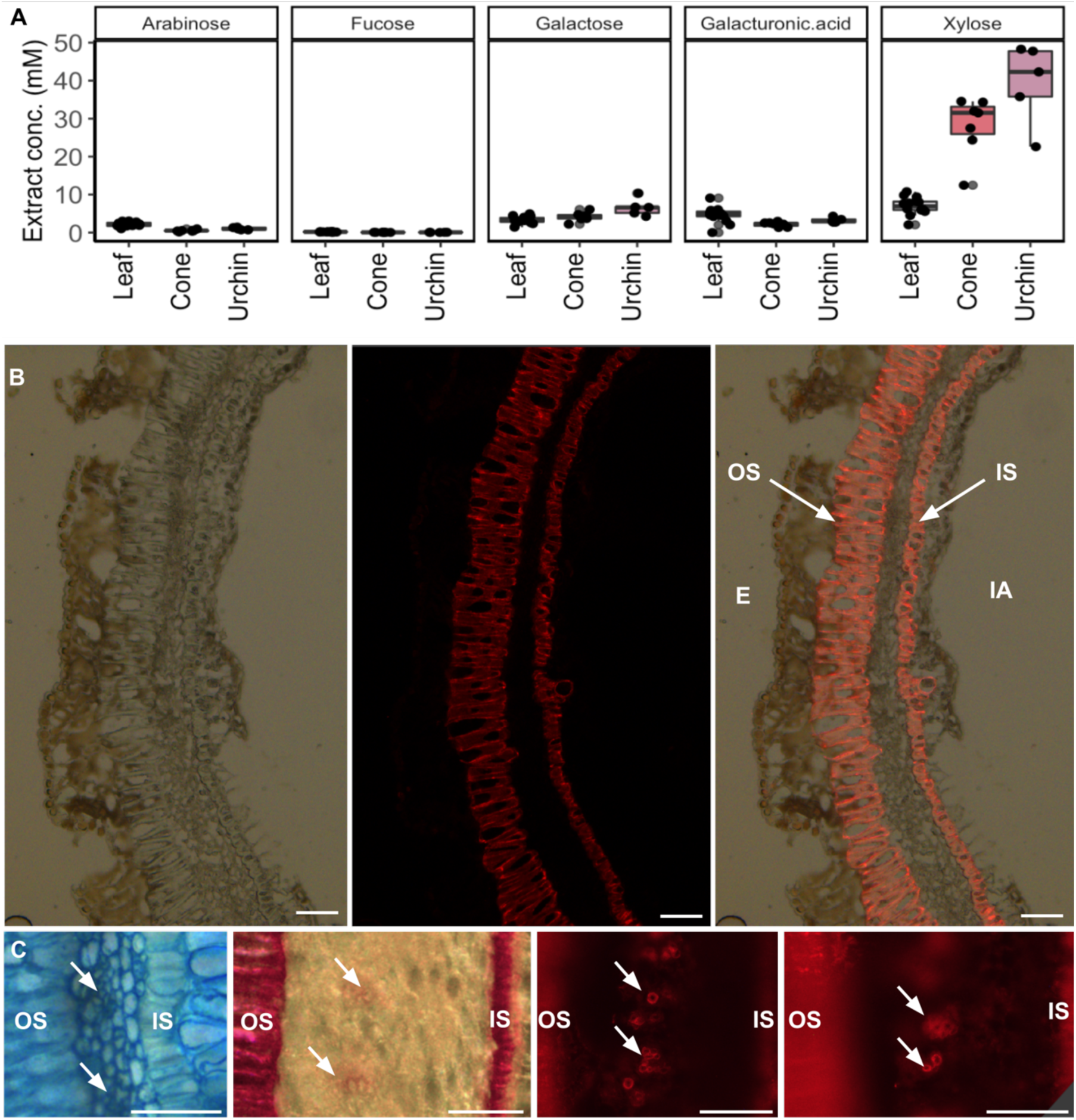
Composition of gall cell walls are altered to be highly enriched in xylan. A: Concentration of five sugars in cell wall residue hydrolysate. Glucose potentially derived from cell wall polymers cannot be accurately measured due to starch contamination. B: LM10 immunofluorescence staining signal for xylan. Left: differential interference contrast transmitted light. Center: Alexa-fluor 647 secondary antibody conjugated to LM10 primary antibody. Right: overlay. OS: outer sclerenchyma, IS: inner sclerenchyma, E: exterior, IA: interior airspace. Scale bar = 50 µm. C: Views of vascular bundles in sponge layer, from left to right: Toluidine Blue O, Weisner stain, LM10, LM10. In each case the outer sclerenchyma cells are shown on the left, sponge layer containing vascular bundles (arrows) in the middle, and inner sclerenchyma on the right (mostly cropped out in LM10 images due to focus and saturation issues). All scale bars = 50 µm, uncropped source images available in Supplemental Figure 8.

The extraordinarily high levels of xylose suggest enrichment of a polymer composed largely of xylose in gall tissue. One natural candidate is xylan, which is named after and usually enriched in xylem tissue and other vasculature fibers (57, 58). The antibody LM10 selectively binds to and is used to detect xylan. We performed immunofluorescence microscopy with LM10 raised in mice as primary antibodies and anti-mouse IgG conjugated to Alexa-fluor 647 as a secondary antibody as shown in Figure 4B. This revealed high concentrations of xylan in the same two sclerenchyma cell layers which are highly lignified (Figure 3A, B). The colocalization of xylan and lignin deposition is suggestive of a mechanical defense role for these two cell layers. Furthermore, at higher magnification and exposure there were bundles of cells present in the sponge layer between these two, colocalizing with the lignified bundles revealed by Weisner stain (arrows in Figure 3). These bundles as viewed with toluidine blue O stain, Weisner stain, and two views of LM10 immunostain are shown in orientation-matched views in Figure 4C (uncropped source images available in Supplemental Figure 11). The colocalization of lignin and xylan in this particular bundled spatial pattern strongly suggests these are the vascular bundles of the gall, and their less-consistent organization compared to normal vascular bundles likely reflects imperfect control of plant developmental morphology on part of the gall-inducing wasps, as noted previously regarding the alteration of existing vasculature in galls (59). The cell-layer specific alteration of lignin and polysaccharide composition – the two primary constituents of plant cell walls – indicates that galling insects exert a large degree of control over plant growth and metabolism in the development of galls.

## Discussion

We leveraged metabolomics, three-dimensional light microscopy, lignin composition analysis, histology, and immunomicroscopy to study the biology of galls. In doing so, we revealed many similarities and several key differences in the metabolic and morphological changes associated with the gall-induction process between two types of gall-inducing wasps. We observe dramatic alteration in metabolite composition in two gall types produced from the same tissue of the same host. While many of the changes to the metabolome are consistent across both gall types, some such as abscisic acid and hexose phosphates are strikingly different. The metabolites with consistent increases in concentration (Dataset S7) are candidates for the shared induction mechanism of galls, whereas those with different concentration changes in the two gall types may be responsible for the specific gall morphology. We have further demonstrated that the cell wall lignin and polysaccharide composition of galls differs substantially from the normal plant tissue from which they arise.

We also present several lines of evidence for *de novo* vascularization in cone galls, a surprising finding given the leaf tissue from which the galls derive is terminally differentiated. It has long been known that gall-inducing insects modify and enlarge existing vasculature to deliver nutrients to the gall (28, 53, 54). Galls induced by *Agrobacterium* were long thought to lack vasculature (60) but eventually shown to contain a vascular system organized somewhat differently than that found in normal plant tissue (61, 62). Our finding of *de novo* vascularization in insect-induced galls suggests a similar slow-discovery process may be at play for insect-induced galls. Leafy galls induced by *Rhodococcus fascians* have also been shown to induce neovascularization (63), so while our report is to our knowledge the first describing neovascularization in insect-induced galls, neovascularization is widely accepted to occur in some galls, specifically bacteria-induced galls.

This detailed analysis of the morphological, metabolic, and structural changes found in cynipid galls invites comparison to better understood galls such as the crown gall induced by *Agrobacterium*. The key principle of crown gall induction by *Agrobacterium* is transfer and expression of a relatively short stretch of ‘T-DNA’ that comprises part of the tumor-inducing plasmid (64). This stretch of DNA encodes enzymes in the biosynthetic pathway for auxin (65) and cytokinin (66), which result in altered phytohormone levels and ratios in crown gall (67). The mechanism of gall induction in root knot nematodes is less well understood, but it is notable that the gall-inducing nematodes have been shown to synthesize auxin (68), which suggests synthesis of plant hormones is a common strategy to manipulate plant tissue to expand. Cynipid galls are much more morphologically complex than either of these better-characterized systems, and there is much more diversity in gall size, shape, location, and color. This diversity suggests that the mechanics of gall induction vary between different cynipid wasps, which is supported by our data demonstrating different changes to phytohormones. Nonetheless, the phylogenetic distribution of the galling habit within cynipid wasps suggests it is ancestral, and therefore at least some of the core mechanics are likely to be conserved (32).

The complex and colorful structures of galls have captured the imagination of naturalists for millennia, and demonstrate a mastery of inter-kingdom manipulation that remains unparalleled by current plant molecular biologists. Many practices used to modify and manipulate plants are still reliant on the same techniques adapted from natural plant engineers (i.e., *Agrobacteria*) several decades ago becoming the foundation of plant genetic transformations. Thus, looking for more examples in nature of non-model, non-traditional systems to expand our perspective on the degree to which plants can be reprogrammed may inspire novel approaches to engineering plants in general. Elucidating the molecular basis of the induction of complex galls may provide the blueprint to redefining the landscape to redesigning entirely new cellular, morphological, and physiological architectures in plants.

## Materials and Methods

### Gall collection

We monitored an arboretum collection of approximately one hundred species of oak trees for galls from spring to autumn. Dozens of gall types were found, of which two types of galls proved to be relatively abundant: the cone gall induced by *Andricus kingi* and the urchin gall induced by *Antron douglasii*, both on the valley oak *Quercus lobata*. Both of these galls were found on the abaxial and adaxial surfaces of leaves between June and August of 2019-2022, with the cone galls being more abundant and appearing somewhat earlier. Both were markedly concentrated in particular trees; one valley oak would often contain hundreds of galls while none could be seen on other valley oaks only a dozen meters away (Supplemental Figure 1). Furthermore, the cone galls in particular were found to cluster on particular branches -it was common to see one branch supporting many times more galls per leaf than an adjacent branch, a somewhat surprising finding given that the gall-inducing insects can fly. Galls were collected in the UC Davis arboretum (38°31”46.0”N 121°45”45.3’W) and Putah Creek Riparian Reserve (38°31’19.7”N 121°46”50.1”W) between May 2019 and September 2020. Initially, all galls identified on oak trees were collected, making use of the 89 species of oak trees in the Peter J Shields collection. Over 20 types of galls were initially collected. The cone gall and urchin gall induced by *Andricus kingii* and *Antron douglasii* on valley oak (*Quercus lobata)* were selected on the basis of morphological complexity and abundance in the study area for further collection and analysis, and 1000-2000 galls of these two species were gathered, at times individually divided into classes on the basis of mass / growth stage, at times in mass collections for large-scale metabolite analysis. Galls were removed from the tree and flash frozen in liquid nitrogen as quickly as possible. Date of collection and specific tree of origin were noted for each gall sample (Dataset S1, Supplemental Figure 1).

### Laser Ablation Tomography

Fresh gall samples were sent to LATscan (State College, PA) to perform laser ablation tomography. In brief, samples are attached to a piece of pasta as a sacrificial supporting structure, then mounted in the beam path of a microscope from the front and a high-power flat-beam laser from the side. Rapid alternation of microscope image captures and laser pulses allows for rapid acquisition of several thousand serial ‘slice’ images through the entire sample.

### Metabolite extraction

Metabolites were extracted using a protocol adapted from (69). Galls and leaves were flash-frozen in liquid nitrogen and stored at −80 °C until processing. Samples were lyophilized, then disrupted with a steel ball in a ball mill at 30 Hz for 20 minutes, yielding a fine powder. Powder was weighed, then 80 µL of methanol was added per mg. Samples were vortexed for 1 minute, then incubated at room temperature for 20 minutes with continuous mixing, centrifuged at 20,000 g for 5 minutes and the supernatant filtered through 0.45 µm PTFE filters.

### Mass spectrometry

In preparation for LC-MS analysis, oak gall extracts were first dried in a SpeedVac (SPD111V, Thermo Scientific, Waltham, MA), then resuspended in 100% MeOH containing an internal standard mix of isotopically labeled compounds (∼15 µM average of 5-50 µM of 13C,15N Cell Free Amino Acid Mixture, #767964, Sigma; 10 µg/mL 13C-trehalose, #TRE-002, Omicron; 10 µg/mL 13C-mannitol, ALD-030, Omicron; 2 µg/mL 13C-15N-uracil, CNLM-3917, CIL; 5.5 µg/mL 15N-inosine, NLM-4264, CIL; 4 µg/mL 15N-adenine, NLM-6924, CIL; 3 µg/mL 15N-hypoxanthine, NLM-8500, CIL; 5 µg/mL 13C-15N-cytosine, #294108, Sigma; 2.5 µg/mL 13C-15N-thymine, CNLM-6945, CIL;, 1 µg/mL 2-amino-3-bromo-5-methylbenzoic acid, R435902, Sigma), with resuspension volume of each varied to normalize by biomass for each sample group.

UHPLC normal phase chromatography was performed using an Agilent 1290 LC stack, with MS and MS/MS data collected using a QExactive HF Orbitrap MS (Thermo Scientific, San Jose, CA). Full MS spectra was collected from m/z 70 to 1050 at 60k resolution in both positive and negative ionization mode, with MS/MS fragmentation data acquired using stepped then averaged 10, 20 and 40 eV collision energies at 15,000 resolution. Mass spectrometer source settings included a sheath gas flow rate of 55 (au), auxiliary gas flow of 20 (au), spray voltage of 3 kV (for both positive and negative ionization modes), and capillary temperature or 400 degrees C. Normal phase chromatography was performed using a HILIC column (InfinityLab Poroshell 120 HILIC-Z, 2.1 × 150 mm, 2.7 µm, Agilent, #683775-924) at a flow rate of 0.45 mL/min with a 3 μL injection volume. To detect metabolites, samples were run on the column at 40 °C equilibrated with 100% buffer B (99.8% 95:5 v/v ACN:H2O and 0.2% acetic acid, w/ 5 mM ammonium acetate) for 1 minute, diluting buffer B down to 89% with buffer A (99.8% H2O and 0.2% acetic acid, w/ 5 mM ammonium acetate and 5 µM methylene-di-phosphonic acid) over 10 minutes, down to 70% over 4.75 minutes, down to 20% over 0.5 minutes, and isocratic elution for 2.25 minutes, followed by column re-equilibration by returning to 100% B over 0.1 minute and isocratic elution for 3.9 minutes. Samples consisted of 8 biological replicates each and extraction controls, with sample injection order randomized and an injection blank of of 100% MeOH run between each sample.

Metabolite identification was based on exact mass and comparing retention time (RT) and fragmentation spectra to that of standards run using the same LC-MS method. LC-MS data was analyzed using custom Python code (Bowen, B. P. Analysis of Metabolomics Datasets with High-Performance Computing and Metabolite Atlases. 431–442 (2015). doi:10.3390/metabo5030431), with each detected peak assigned a level of confidence, indicated by a score from 0 to 3, in the compound identification. Compounds given a positive identification had matching RT and *m/z* to that of a standard, with detected m/z ≤ 5 ppm or 0.001 Da from theoretical as well as RT ≤ 0.5 minutes. A compound with the highest level of positive identification (score of 3) also had matching MS/MS fragmentation spectra. An identification was invalidated when MS/MS fragmentation spectra collected for the feature did not match that of the standard.

### Molecular networking

The LC-MS files were run via MZmine2 version 2.39 workflow to generate a list of features, which were putatively annotated using the Global Natural Products Social Molecular Networking (GNPS) tool (70). This pipeline produced molecular networking files for positive (13918 features) and negative polarities (13562). Filtering accepted features with retention time > 0.6 min (post solvent front), maximum peak height > 1e6, and max peak height fold-change between sample and extraction control > 10, resulting in 8690 and 6305 features in negative and positive mode, respectively. The filtered features were merged into a single molecular network (14995 nodes) created in Cytoscape software version 3.9.1 (70, 71) following step-by-step procedure (72). The average peak height in Leaf control (n=16), Urchin (n=32) and Cone (n=40) galls was calculated and painted on each node as pie charts. This was followed by fold-change calculation between average peak height of Urchin or Cone divided by Leaf value; +1 was added to both numerator and denominator to avoid erroneous division by 0. Annotations with cosine score (MQScore) match to library compounds > 0.7 were (1886 nodes) labeled in the networks. NPClassifier was used to determine metabolite classifications of the annotations (33).

Molecular network of combined HILIC untargeted metabolomics without cosine thresholding: https://www.ndexbio.org/viewer/networks/02f90a6c-dafd-11ed-b4a3-005056ae23aa

Molecular network of combined HILIC untargeted metabolomics cosine threshold 0.7 organized by mass feature cosine score: https://www.ndexbio.org/viewer/networks/0d278c0e-dafd-11ed-b4a3-005056ae23aa

Molecular network of combined HILIC untargeted metabolomics cosine threshold 0.7 organized by NP Classifier class: https://www.ndexbio.org/viewer/networks/133ba370-dafd-11ed-b4a3-005056ae23aa

### Lignin quantification

Lignin content was measured using the thioglycolic acid (TGA) method following Suzuki *et al* 2009 (73). 1 mL 3N HCl and 0.1 mL TGA were added to 15 mg of biomass. Samples were then incubated at 80°C for 3 hours, centrifuged for 10 minutes at 16,100 g and the supernatant discarded. 1 mL sterile water was added to the pellet, and vortexed for 30 seconds, and the sample was again centrifuged with the same conditions. 1 mL 1N NaOH was added to the pellet and the sample was allowed to shake at 80 rpm at room temperature for 16 hours, then centrifuged with the same conditions. 1 mL supernatant was transferred to a new tube and 0.2 mL of 12N HCl was added in a fume hood. The samples were then incubated at 4°C for 4 hours and centrifuged 10 minutes at 16,100 g. The supernatant was discarded and the pellet was dissolved in 1 mL 1N NaOH. Dilutions prepared in 1N NaOH were used to measure absorbance (A280). Lignin concentrations were compared with Wilcoxon rank sum test using the Benjamini-Hochberg method for adjustment for multiple comparisons.

### Alcohol-insoluble residue (AIR) preparation

AIR prep was adapted from (74). AIR extracts were prepared by adding ∼15 mg of flash-frozen tissue to 1 mL 100% EtOH. The tissue was then ground in a ball mill at 20 Hz for 5 minutes, heated at 100°C for 30 minutes with periodic shaking, cooled to room temperature and centrifuged at 21,000 g for 5 minutes. The supernatant was discarded and 1 mL 70% EtOH added and vortexed, then centrifuged at 20,000 g for 1 minute. These three steps were repeated until the supernatant was clear, and that clear supernatant discarded. 1 mL of acetone was then added and the samples vortexed, centrifuged at 20,000 g for 5 minutes, supernatant discarded, and the samples dried in a speed-vac overnight. The result was a fine powder which was stored at 4°C. .

### Lignin monomeric composition

A small amount (∼1 mg) of AIR extract was loaded into a quartz tube for Pyro-GC MS analysis using the methodology adapted from (75). Pyrolysis of biomass was performed with a Pyroprobe 5200 (CDS Analytical Inc., Oxford, PA, USA) connected with GC/MS (Thermo Electron Corporation with Trace GC Ultra and Polaris-Q MS) equipped with an Agilent HP-5MS column (30 m x 0.25 mm inner diameter, 0.25 µm film thickness). The pyrolysis was carried out at 650 °C. The chromatograph was programmed from 50 °C (1 min) to 300 °C at a rate of 20 °C/min; the final temperature was held for 10 min. Helium was used as the carrier gas at a constant flow rate of 1 mL/min. The mass spectrometer was operated in scan mode and the ion source was maintained at 300 °C. The compounds were identified by comparing their mass spectra with those of the NIST library. Peak molar areas were calculated for the lignin degradation products, and the summed areas were normalized.

### Trifluoroacetic acid (TFA) hydrolysis

TFA hydrolysis and high-pressure anion exchange chromatography (HPAEC) was adapted from (76). 5-10 mg of AIR was transferred to a new tube using a +-0.01 mg scale to record transferred mass. 1 mL 2M TFA was added to each sample in a screw-top tube and vortexed. Samples were heated to 120°C for 1 hour, vortexing for 10 seconds every 15 minutes. After cooling to room temperature, samples were centrifuged at 20,000 g for 1 minute, as much supernatant as possible discarded, and the remainder removed by speed-vac overnight. The dried pellet was dissolved in 1 mL water and shaken at 1000 rpm 30°C for 1 hour, then filtered through 0.45 μm nitrocellulose filters. Samples were then diluted in water for HPAEC coupled with pulsed amperometric detection. As described in text, several dilution ratios were ultimately required, ranging from 1/10 to 1/640. NaOH was used as needed to bring all samples within the range of 4-9 pH.

### Microscopy

Samples were either kept at 4°C and imaged within 1 week of collection or flash frozen in liquid nitrogen and stored at −80°C. Sectioning was performed with a vibratome to generate ∼50 μm sections or with a cryotome to generate ∼12 μm sections. While several methods of sample fixation were performed, the best results were achieved with unfixed samples embedded in 7% agarose for vibratome sectioning or “Optimal cutting temperature” (Sakura Tissue-Tek OCT, part number 4583) cryotomy embedding fluid. For each stain, several concentrations and staining periods were attempted, and the most informative selected for further work. Imaging was performed with a fluorescent microscope also equipped with an RGB camera, all images except for the immunomicroscopy are real-color, with white-balance adjusted as well as possible to match printed images to the image in the eyepiece.

### Data analysis

Data analysis was performed with Rstudio (Version 2022.07.0+548 macOS), primarily using the Tidyverse package for data manipulation and ggplot2 for visualization. Figures were assembled with Google Drawings.

## Data availability/ Supplementary information

All data produced in this project are available in the main Figures, Supplemental Figures, Supplemental Datasets, and Supplemental Movies.

## Acknowledgements

This work was supported by the UC Davis Katherine Esau Junior Faculty Fellowship to P.M.S. This work was also conducted as part of the DOE Joint BioEnergy Institute (http://www.jbei.org) supported by the U. S. Department of Energy, Office of Science, Office of Biological and Environmental Research, through contract DE-AC02-05CH11231 between Lawrence Berkeley National Laboratory and the U. S. Department of Energy. We thank Denise Snichnes and Steve Ruzin at the UC Berkeley RCNR Berkeley Biological Imaging Facility for advice and assistance with microscopy. We thank LATscan for LAT imaging, Universitas 21 (U21) for funding of LAT. We thank Neelima Sinha and Lynn Kimsey for useful discussions and feedback. The work (proposal:505892 DOI:10.46936/10.25585/60000461)) conducted by the U.S. Department of Energy Joint Genome Institute (https://ror.org/04xm1d337), a DOE Office of Science User Facility, is supported by the Office of Science of the U.S. Department of Energy operated under Contract No. DE-AC02-05CH11231.

## Author Contributions

Designed research: K.M.; P.M.S. Performed research: K.M.; Y.T.; Y.C.C.; K.B.L. Analyzed data: K.M.; V.N.; B.P.B.; Y.T.; Y.C.C.; S.S.; A.Z. Wrote the paper: K.M.; P.M.S.

Advised on project: T.R.N.; A.E.; H.V.S.; P.M.S.

## Supplemental Figures

**Supplemental Figure 1:**
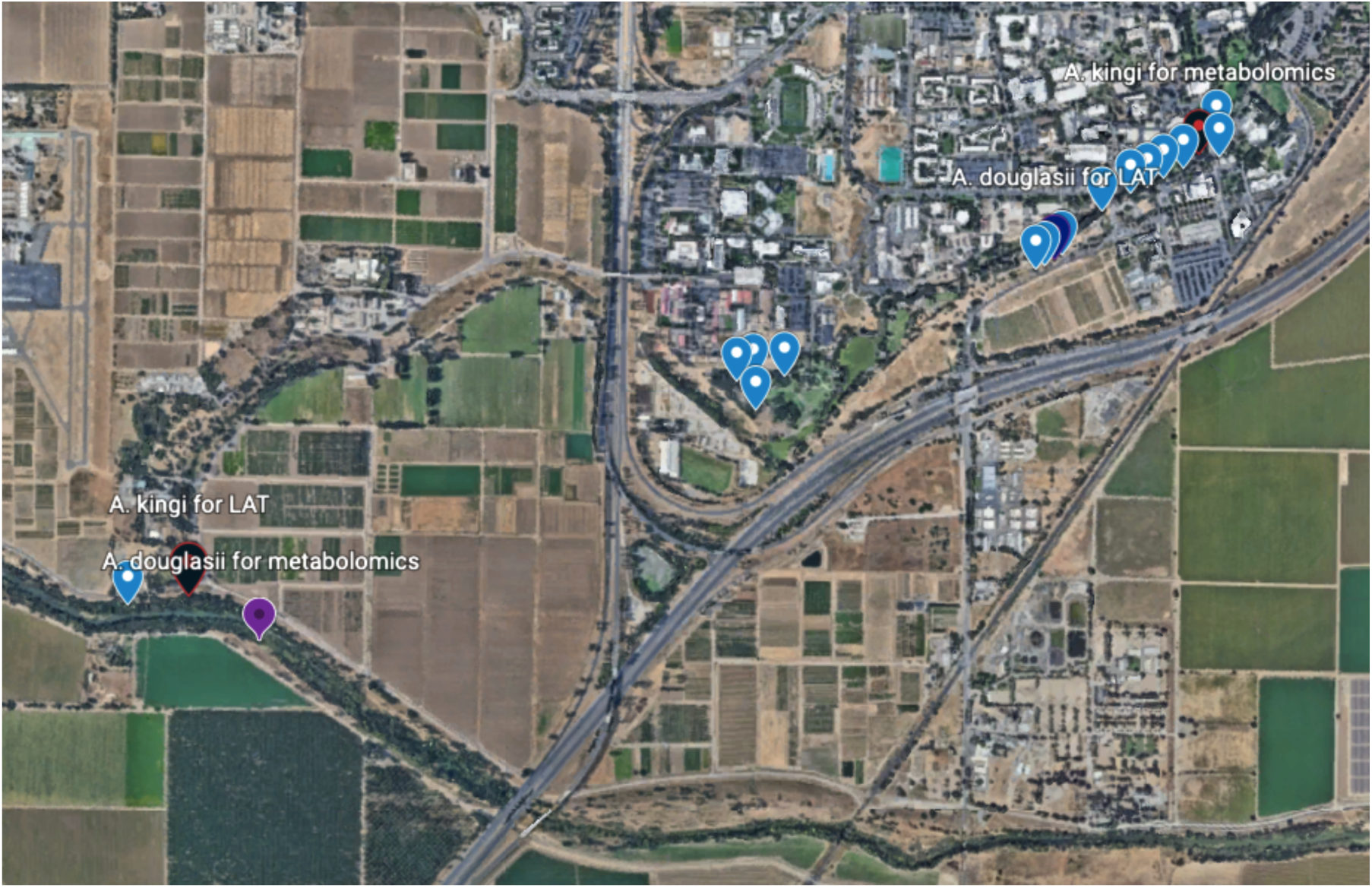
Map of gall collection sites. Round tags indicate individual oak trees from which galls were harvested, red tags indicate harvests of galls induced by *Andricus kingi* that were used in further experiments, purple tags indicate harvests of galls induced by *Antrol douglasii* in further experiments. For each experiment, all galls of a particular type were harvested from one tree on one day.

**Supplemental Figure 2:**
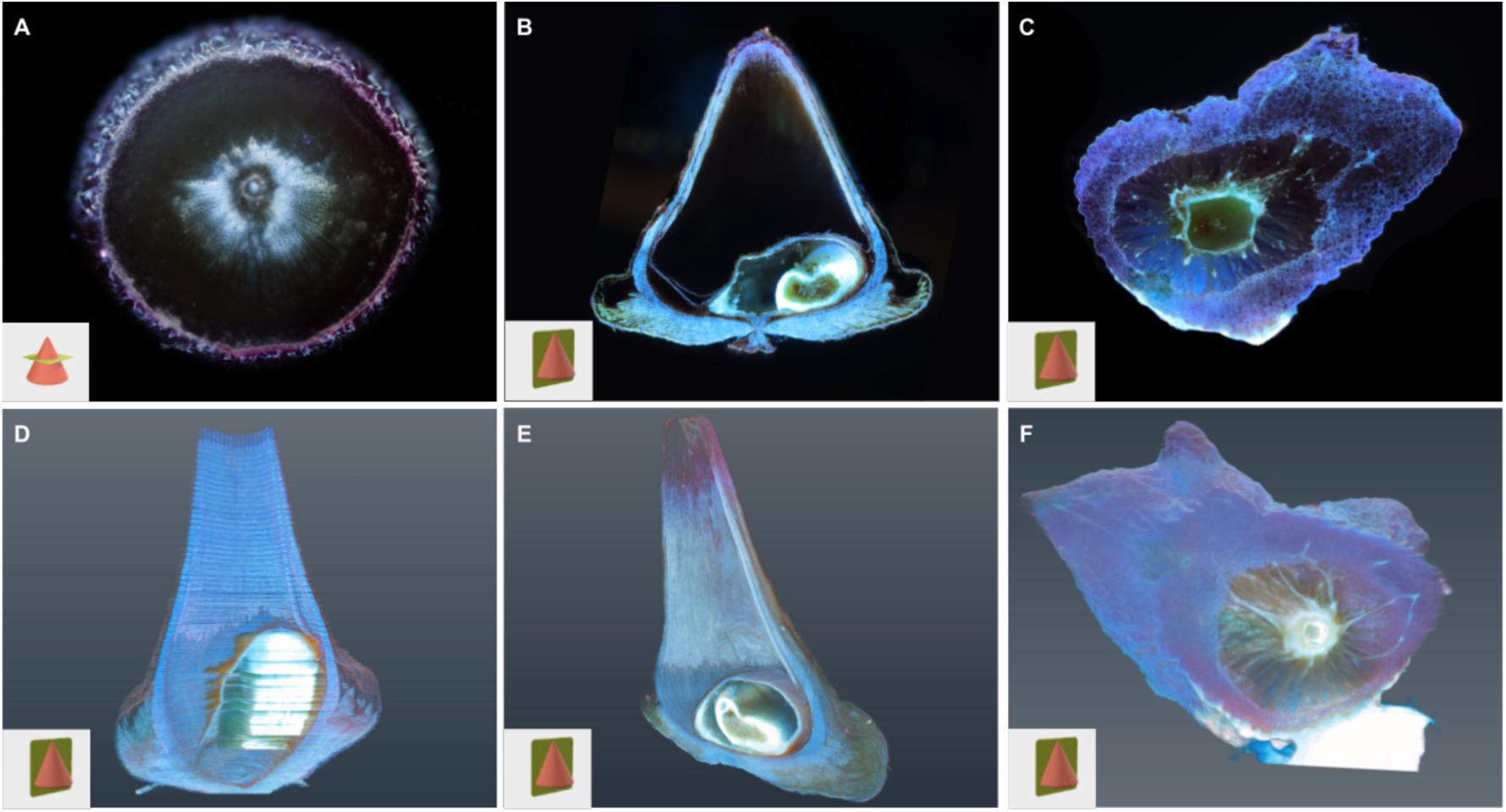
Laser ablation tomography sections and three-dimensional models. A: Transverse section of cone gall. B: Longitudinal sectional of cone gall. C: longitudinal section of urchin gall. D: three-dimensional model of cone gall imaged in transverse sections (apparent horizontal lines are an artifact of aliasing in assembling individual slices and stochastic variance in image brightness), corresponds to supplemental movie 1. E: three-dimensional model of cone gall imaged in longitudinal sections, corresponds to supplemental movie 2. F: three-dimensional model of urchin gall, corresponds to supplemental movie 3. Inset diagrams in lower left corner indicate orientation as viewed, which corresponds to view as imaged with the exception of D, where view shown here is orthogonal to the imaging plane.

**Supplemental Figure 3:**
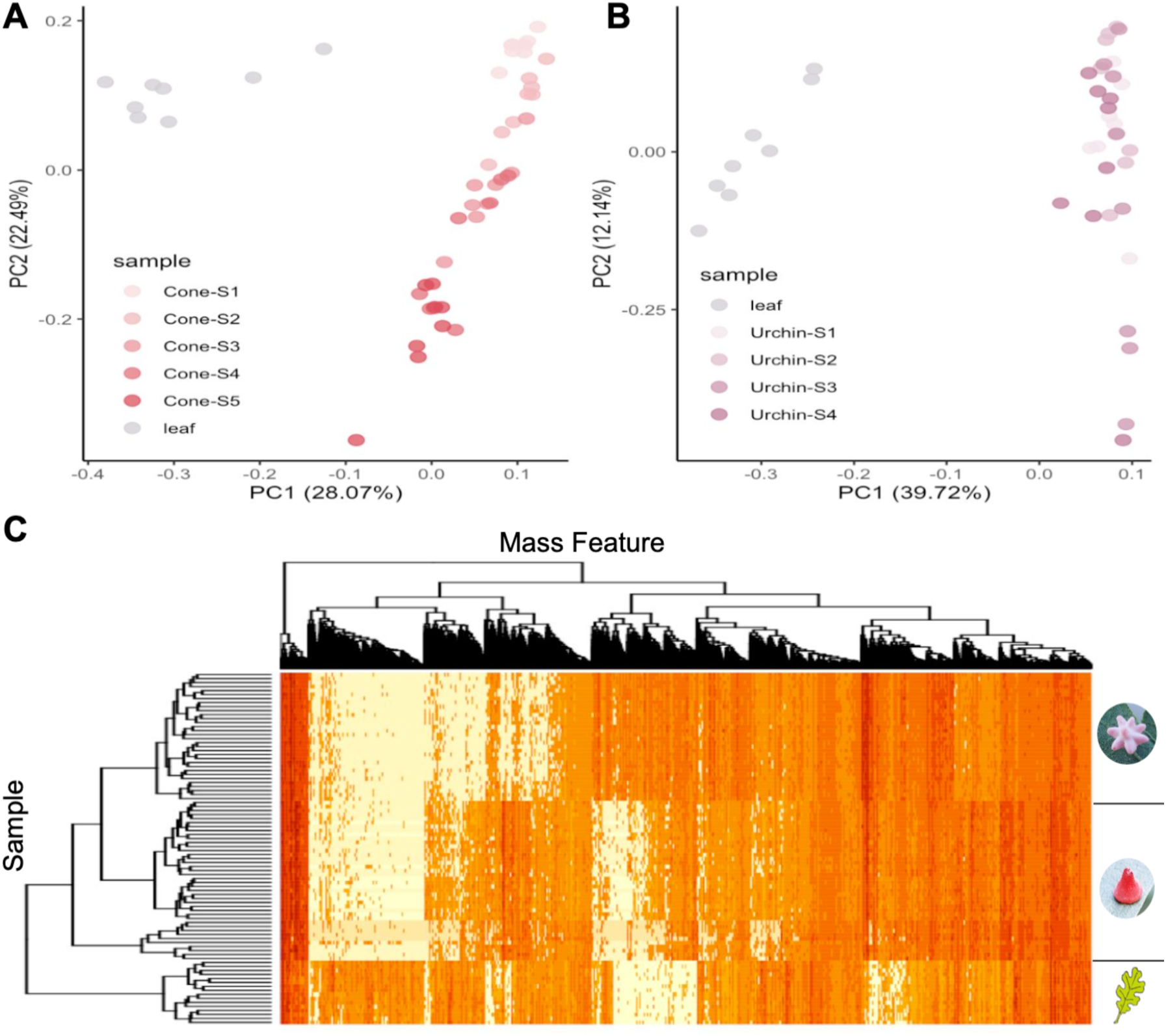
Gall metabolomes cluster by developmental stage. A: PCA of the developmental stages of cone galls. B: PCA of the developmental stages of urchin galls. C: Heatmap of untargeted metabolomics with dendrogram. Rows are samples, columns are metabolites.

**Supplemental Figure 4:**
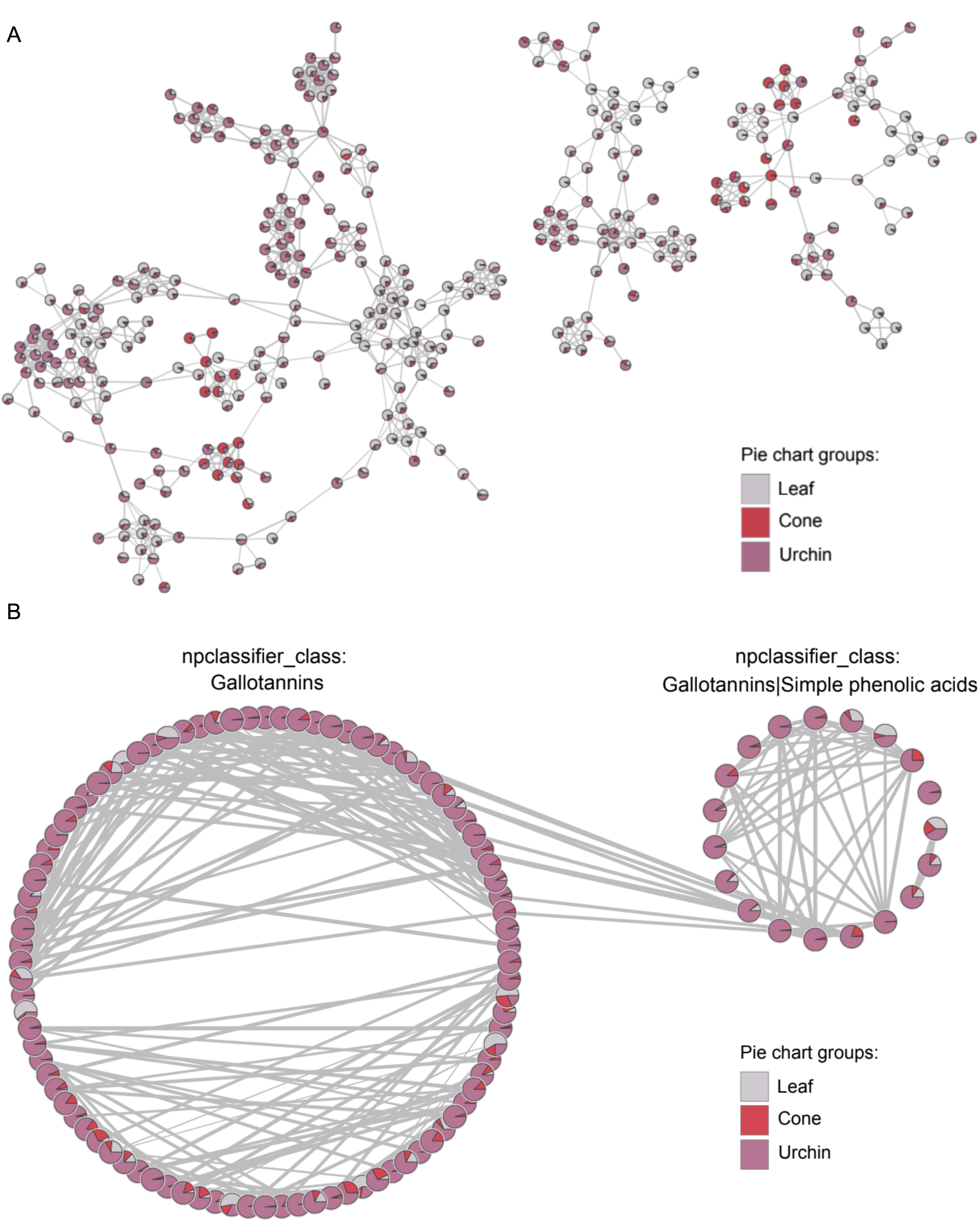
Gall metabolomes are distinct. A: Network from untargeted metabolomics. Interactive networks with putative identifications of mass features available at NDExBio. B: Network of mass features classified as gallotannins by NPClassifier.

**Supplemental Figure 5:**
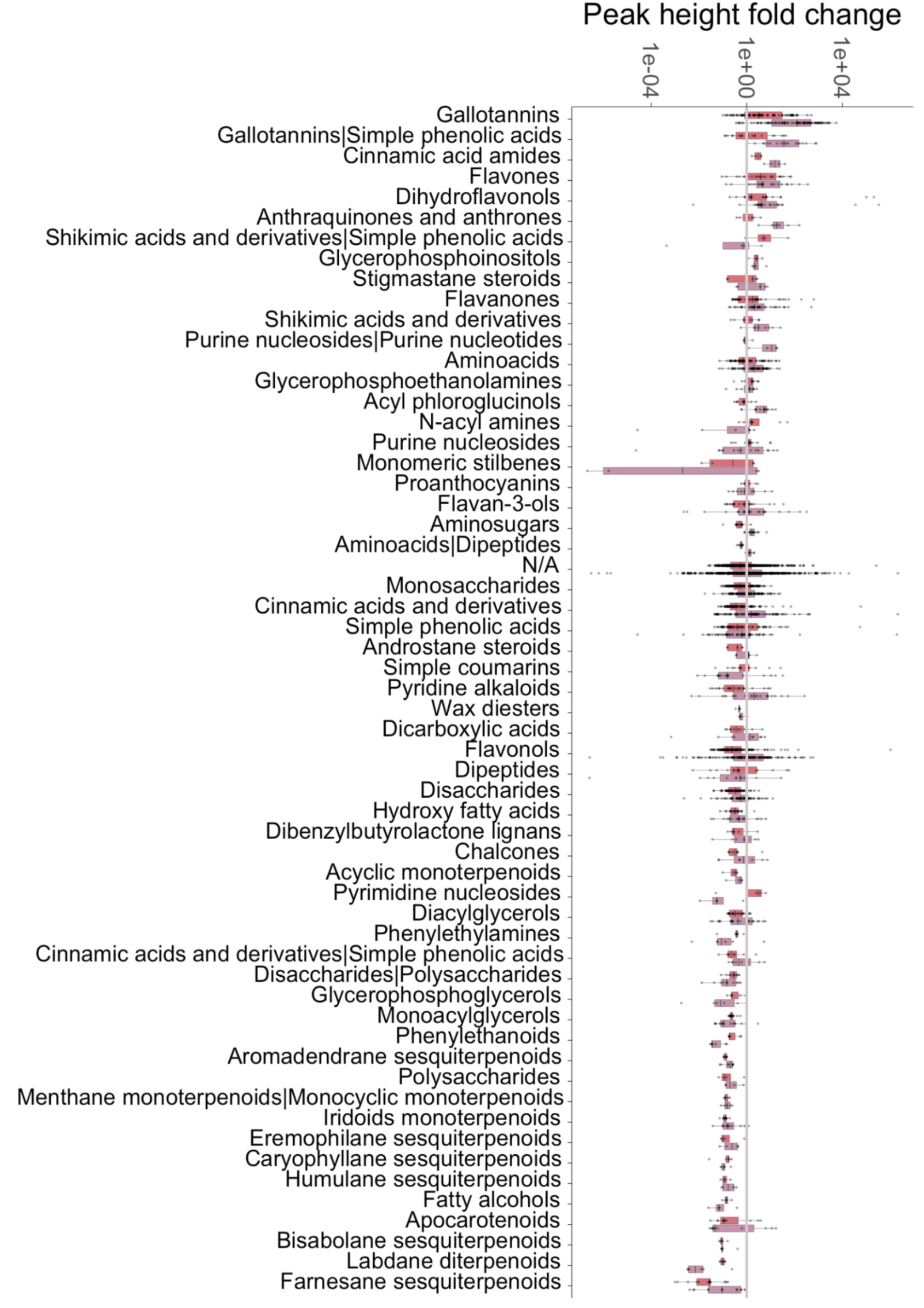
Galls differ in chemical class concentration. Peak height fold change for all NPClassifier categories for both types of gall compared to leaf tissue.

**Supplemental Figure 6:**
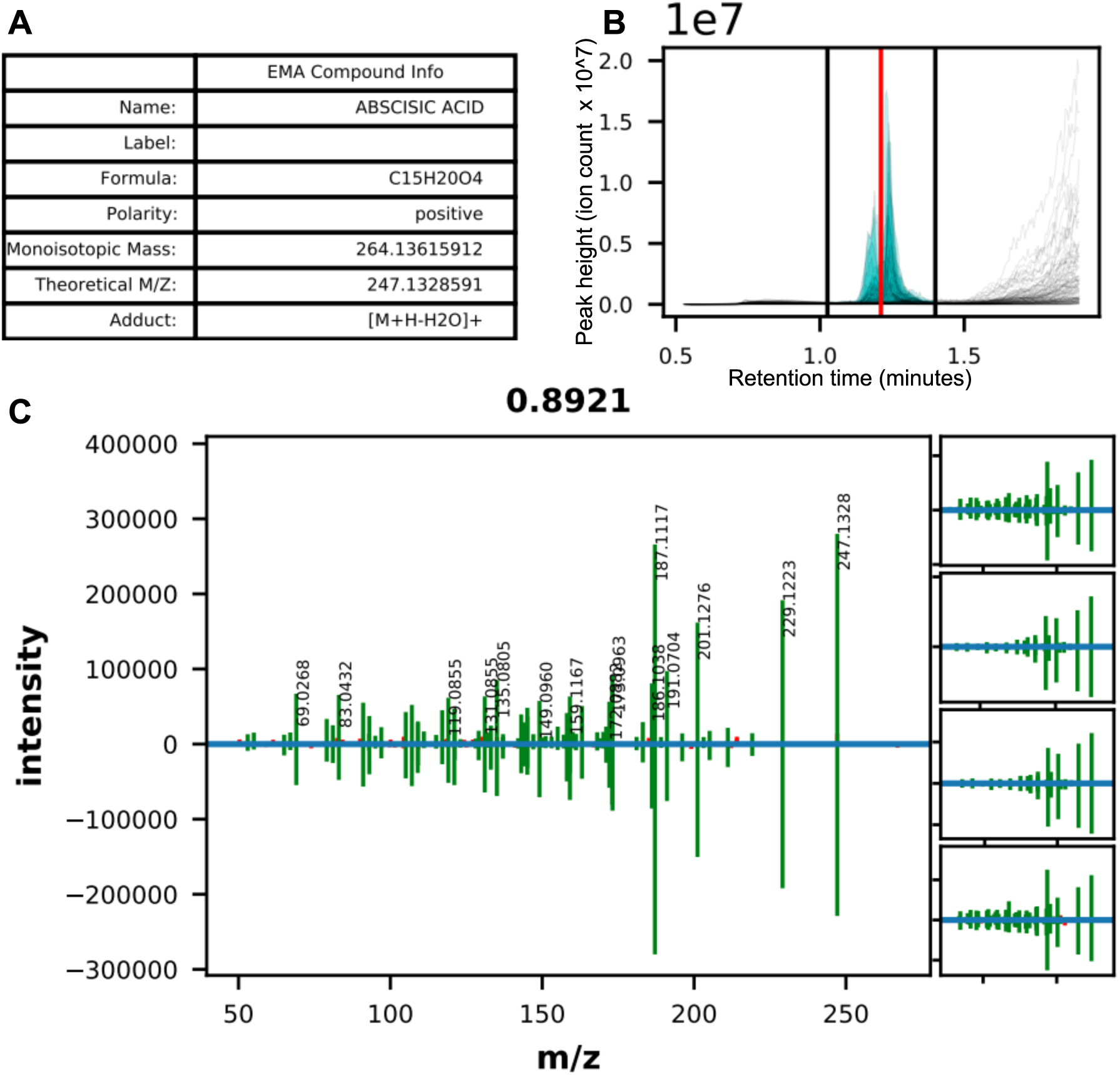
MS-MS details for abscisic acid. A: Information table. B: Extracted ion chromatogram across all files. The predicted retention time is the red vertical line, the black vertical lines are the integration bounds. C: MS-MS mirror plot, our data on top and standard MS-MS fragmentation pattern on bottom. Cosine score = 0.8921. Panels to the right are the next four highest scoring comparisons.

**Supplemental Figure 7:**
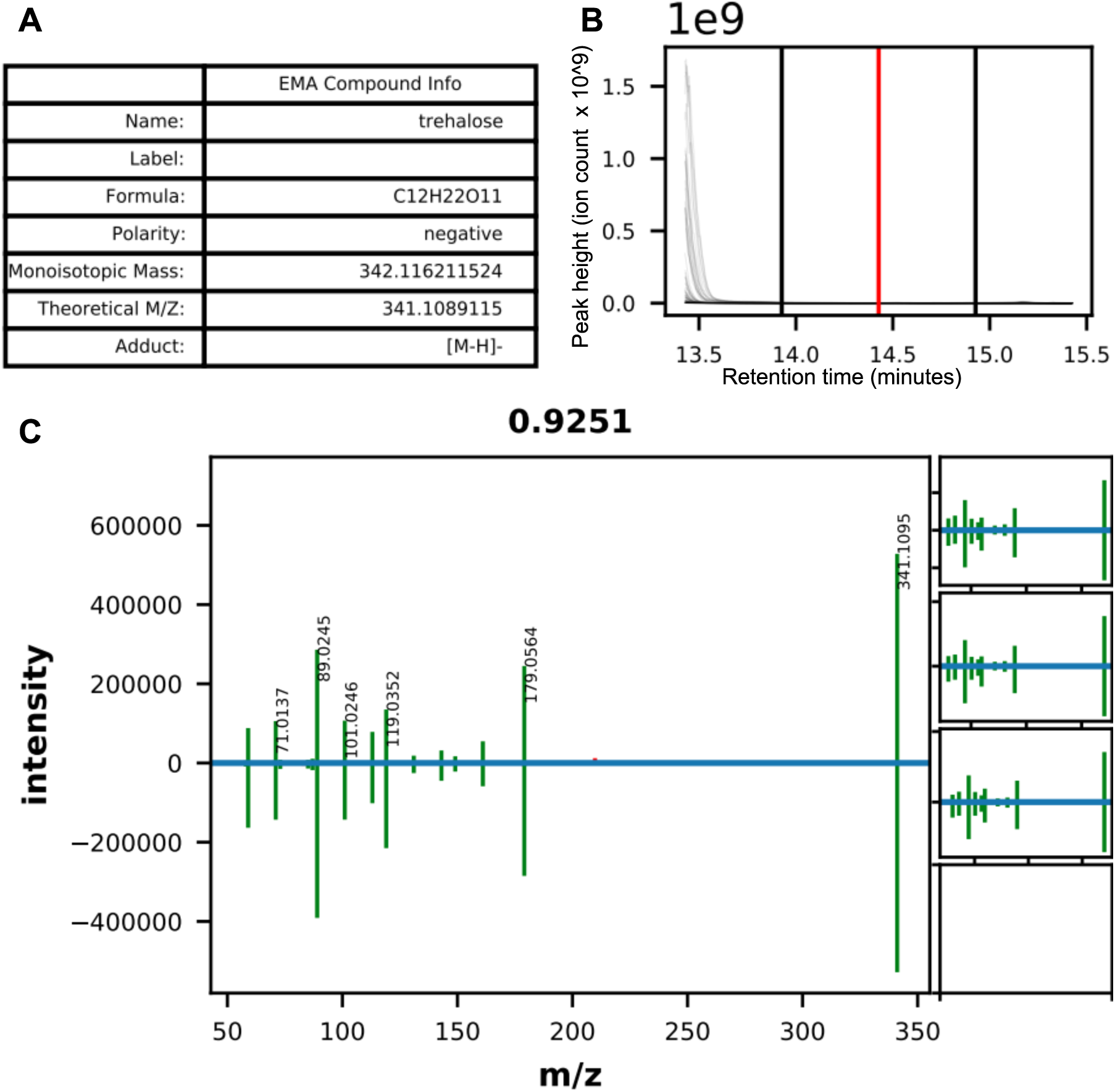
MS-MS details for trehalose. A: Information table. B: Extracted ion chromatogram across all files. The predicted retention time is the red vertical line, the black vertical lines are the integration bounds. Due to software constraints, this graph is too zoomed out on the Y axis because of another peak around 13.2 minutes. The maximum peak intensity for trehalose is 7.65 x 10͞6, which is not visible due to the Y axis scaling. The peak is at 14.42 minutes (very close to the predicted retention time), and the integration bounds are set well, as can be seen in Dataset S5. C: MS-MS mirror plot, our data on top and standard MS-MS fragmentation pattern on bottom. Cosine score = 0.9251. Panels to the right are the next four highest scoring comparisons. In this case, only alternative three matches were found by the software.

**Supplemental Figure 8:**
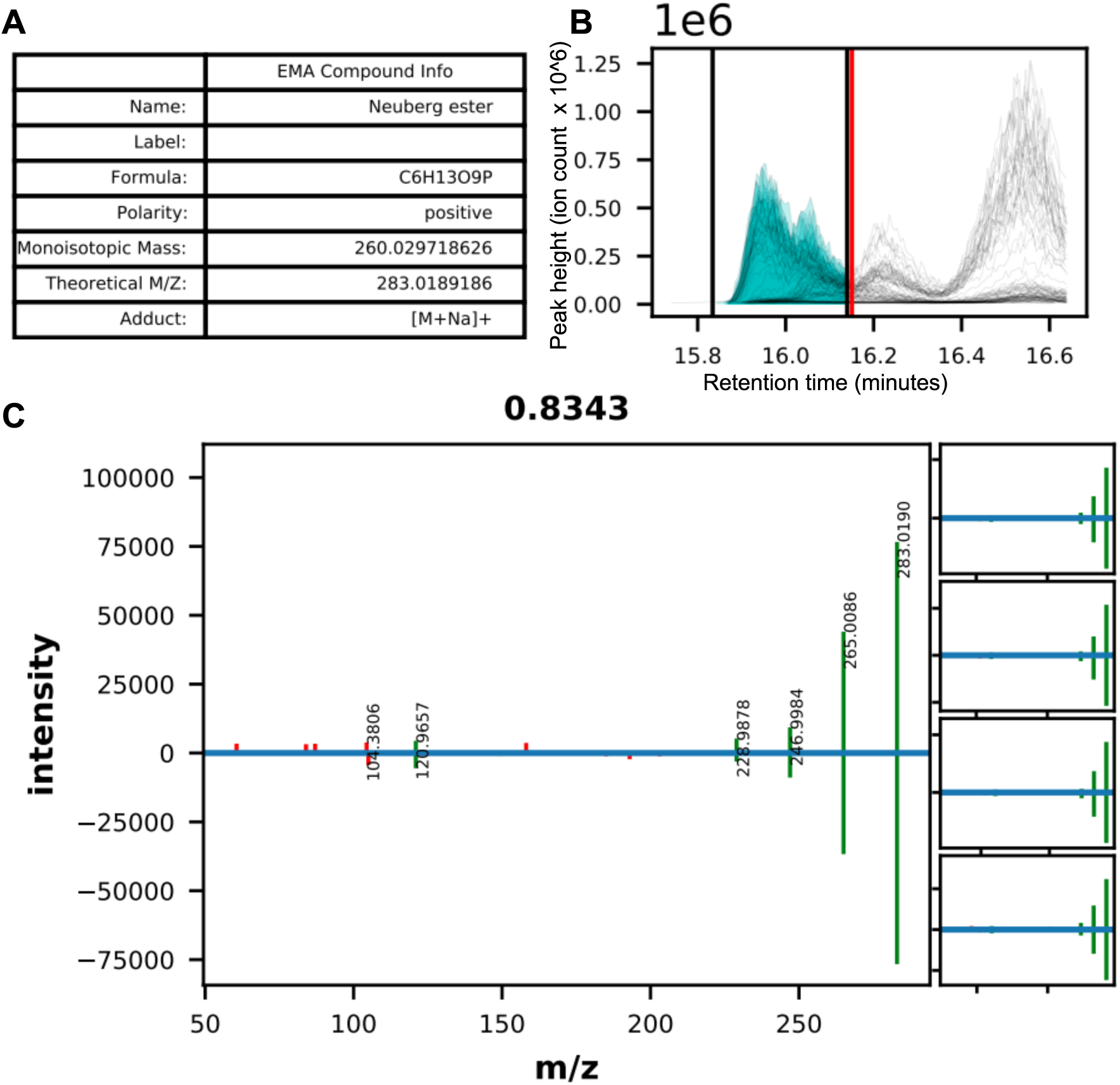
MS-MS details for hexose phosphate. A: Information table. While the software identified this feature as Neuberg ester meaning Fructose-6-phosphate, we are not confident of this specific isomeric identification, only that this is a hexose phosphate. B: Extracted ion chromatogram across all files. The predicted retention time is the red vertical line, the black vertical lines are the integration bounds. C: MS-MS mirror plot, our data on top and standard MS-MS fragmentation pattern on bottom. Cosine score = 0.8343. Panels to the right are the next four highest scoring comparisons.

**Supplemental Figure 9:**
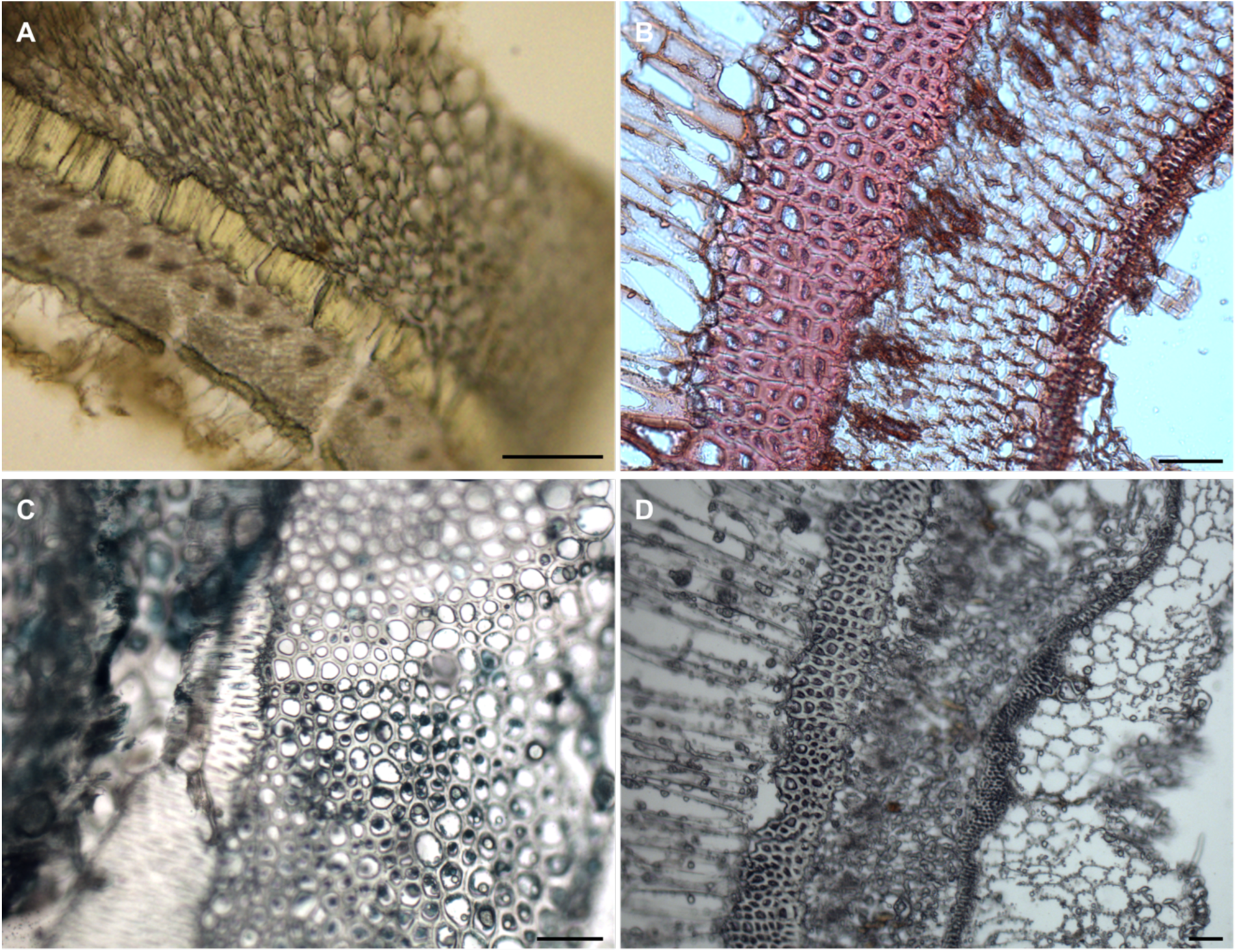
Sample of additional histological stains. A: Cone gall stained with Mäule stain. Scale bar = 100 µm. B: Cone gall stained with Safranin O. Scale bar = 50 µm C: Cone gall stained with FastGreen. Scale bar = 50 µm D: Unstained cone gall. Scale bar = 50 µm

**Supplemental Figure 10:**
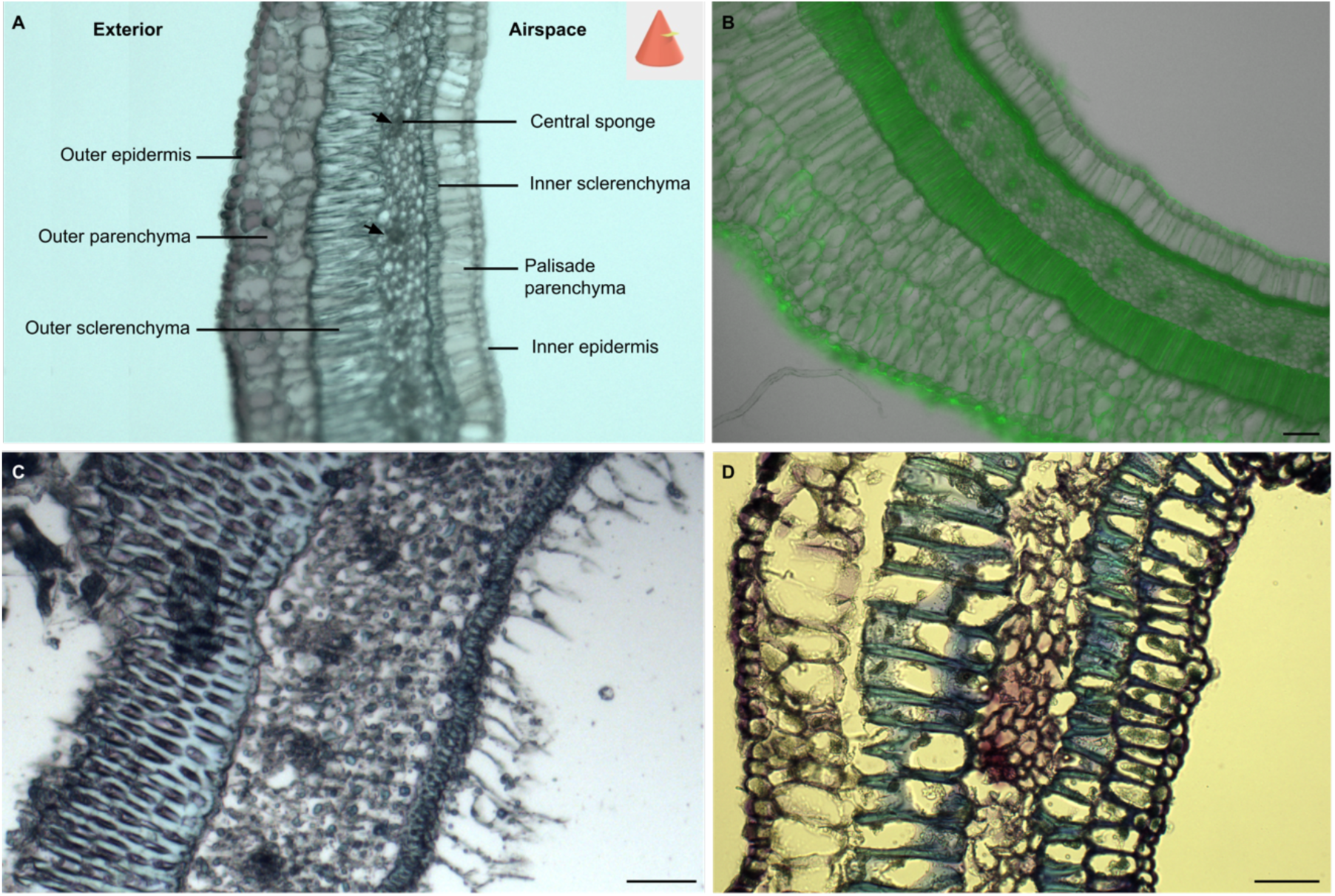
Micrographs of cone galls and map of cell layers. A: Transverse section of cone gall with cell layers labeled. There are 7 distinct cell layers, which can be seen by their morphology and differential uptake of stains. Arrows indicate bundles which are moderately lignified. Outer sclerenchyma and inner sclerenchyma are heavily lignified. B: Transverse section of cone gall imaged by autofluorescence using GFP filter. Scale bar = 50 µm C: 12 µm cryosection stained with toluidine blue O. The thick cell walls of the outer sclerenchyma are stained blue, and the bundles are also clearly visible as dark patches in the central sponge layer. Scale bar = 50 µm. D: 12 µm cryosection stained longer with toluidine blue O. The outer epidermis and parenchyma are stained purple, outer and inner sclerenchyma stained blue, and palisade parenchyma and inner epidermis are stained a dark violet. Scale bar = 50 µm

**Supplemental Figure 11:**
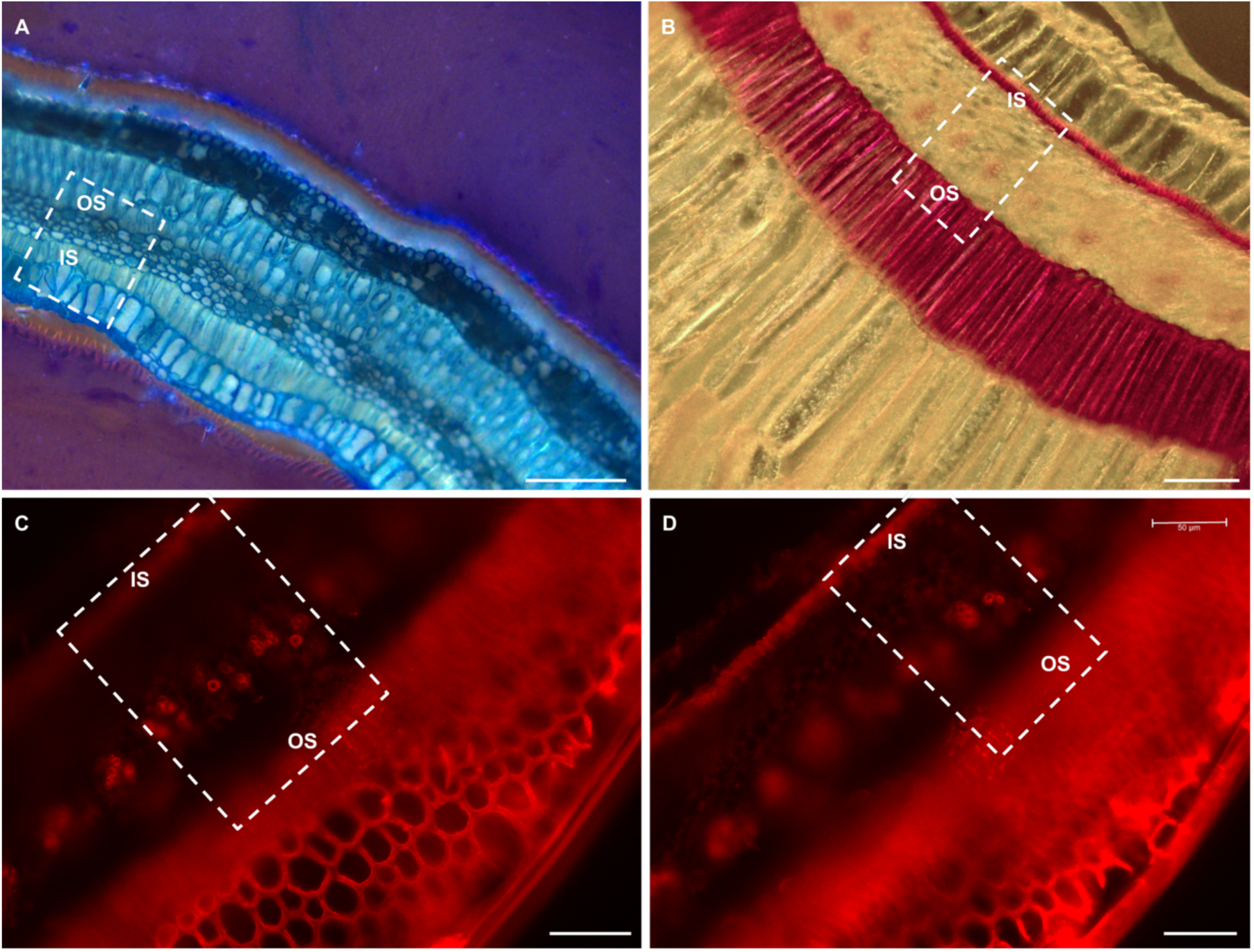
Transverse sections of red cone galls. A: Cone gall stained with toluidine blue O. Scale bar = 100 µm (half-sized bar used to indicate 50 µm in Figure 4C). B: Cone gall stained with Weisner stain. Scale bar = 100 µm (half-sized bar used to indicate 50 µm in Figure 4C). C: Transverse section of cone gall immunostained with LM10. D: Transverse section of cone gall immunostained with LM10. Dashed rectangle goes beyond the edge of the micrograph, explaining the dark gray triangle in bottom right of the rightmost image in Figure 4C. Scale bar = 50 µm. For all micrographs, the dotted rectangle shows the portion of the image used in Figure 4C. OS = outer sclerenchyma, IS = inner sclerenchyma.

## Additional files not included in this document

**Supplemental Movie 1** Cone gall transverse LAT three-dimensional model **Supplemental Movie 2** Cone gall longitudinal LAT three-dimensional model **Supplemental Movie 3** Urchin gall LAT three-dimensional model

**Dataset S1** Dates and locations of gall sampling

**Dataset S2** Positive mode untargeted mass spectrometry dataset

**Dataset S3** Negative mode untargeted mass spectrometry dataset

**Dataset S4** Positive mode targeted mass spectrometry mass feature identification information **Dataset S5** Negative mode targeted mass spectrometry mass feature identification information **Dataset S6** Combined positive and negative mode targeted mass spectrometry dataset **Dataset S7** Metabolites highly enriched in one or both types of gall

**Dataset S8** Lignin subtype analysis pyro gas chromatography mass spectrometry dataset

## References

1. O. Gatjens-Boniche, The mechanism of plant gall induction by insects: revealing clues, facts, and consequences in a cross-kingdom complex interaction. Revista de Biología Tropical 67 (2019).

2. Theophrastus, De Causis Plantarum (London: Heinemann ; Cambridge, Mass. : Harvard University Press, 1976).

3. Theophrastus, Historia Plantarum (1842) (2009).

4. C. Darwin, On the Origin of Species (First Avenue Editions ^TM^, 2018).

5. R. A. Russo, Plant Galls of the Western United States (Princeton University Press, 2021).

6. G. N. Stone, K. Schonrogge, Population and Conservation Ecology, The adaptive significance of insect gall morphology (2003).

7. R. Dawkins, Replicator selection and the extended phenotype. Z. Tierpsychol. 47, 61–76 (1978).

8. D. R. McCalla, M. K. Genthe, W. Hovanitz, Chemical Nature of an Insect Gall Growth-Factor. Plant Physiol. 37, 98–103 (1962).

9. G. Gheysen, M. G. Mitchum, Phytoparasitic Nematode Control of Plant Hormone Pathways. Plant Physiol. 179, 1212–1226 (2019).

10. G. Gheysen, M. G. Mitchum, How nematodes manipulate plant development pathways for infection. Curr. Opin. Plant Biol. 14, 415–421 (2011).

11. D. Veselov, et al., Development of Agrobacterium tumefaciens C58-induced plant tumors and impact on host shoots are controlled by a cascade of jasmonic acid, auxin, cytokinin, ethylene and abscisic acid. Planta 216, 512–522 (2003).

12. T. Yokota, M. Arima, N. Takahashi, Castasterone, a new phytosterol with plant-hormone potency, from chestnut insect gall. Tetrahedron Letters 23, 1275–1278 (1982).

13. H. Yamaguchi, et al., Phytohormones and willow gall induction by a gall-inducing sawfly. New Phytol. 196, 586–595 (2012).

14. H. Suzuki, et al., Biosynthetic pathway of the phytohormone auxin in insects and screening of its inhibitors. Insect Biochem. Mol. Biol. 53, 66–72 (2014).

15. J. F. Tooker, C. M. De Moraes, Feeding by Hessian Fly (Mayetiola destructor [Say]) Larvae on Wheat Increases Levels of Fatty Acids and Indole-3-Acetic Acid but not Hormones Involved in Plant-Defense Signaling. Journal of Plant Growth Regulation 30, 158–165 (2011).

16. J. Traas, Organogenesis at the Shoot Apical Meristem. Plants 8, 6 (2018).

17. Y. Tanaka, K. Okada, T. Asami, Y. Suzuki, Phytohormones in Japanese mugwort gall induction by a gall-inducing gall midge. Biosci. Biotechnol. Biochem. 77, 1942–1948 (2013).

18. L. S. Jankiewicz, M. Guzicka, A. Marasek-Ciolakowska, Anatomy and Ultrastructure of Galls Induced by Neuroterus quercusbaccarum (Hymenoptera: Cynipidae) on Oak Leaves (Quercus robur). Insects 12, 850 (2021).

19. V. C. Martini, A. S. F. Moreira, V. C. Kuster, D. C. Oliveira, Galling insects as phenotype manipulators of cell wall composition during the development of galls induced on leaves of Aspidosperma tomentosum (Apocynaceae). South African Journal of Botany 127, 226–233 (2019).

20. A. T. Formiga, et al., The role of pectic composition of cell walls in the determination of the new shape-functional design in galls of Baccharis reticularia (Asteraceae). Protoplasma 250, 899– 908 (2013).

21. G. N. Stone, J. M. Cook, The structure of cynipid oak galls: patterns in the evolution of an extended phenotype. Proceedings of the Royal Society of London. Series B: Biological Sciences 265, 979–988 (1998).

22. F. Ronquist, J. Liljeblad, Evolution of the gall wasp-host plant association. Evolution 55, 2503– 2522 (2001).

23. L. J. Harper, et al., Cynipid galls: insect-induced modifications of plant development create novel plant organs (2004).

24. J. Hearn, et al., Genomic dissection of an extended phenotype: Oak galling by a cynipid gall wasp. PLoS Genet. 15, e1008398 (2019).

25. E. O. Martinson, J. H. Werren, S. P. Egan, Tissue-specific gene expression shows a cynipid wasp repurposes oak host gene networks to create a complex and novel parasite-specific organ. Molecular Ecology 31, 3228–3240 (2022).

26. D. G. Miller, C. T. Ivey, J. D. Shedd, Support for the microenvironment hypothesis for adaptive value of gall induction in the California gall wasp,Andricus quercuscalifornicus. Entomologia Experimentalis et Applicata 132, 126–133 (2009).

27. B. J. Crespi, D. A. Carmean, T. W. Chapman, Ecology and evolution of galling thrips and their allies. Annu. Rev. Entomol. 42, 51–71 (1997).

28. J. D. Shorthouse, O. Rohfritsch, Biology of Insect-induced Galls (Oxford University Press on Demand, 1992).

29. S. D. Allison, J. C. Schultz, Biochemical responses of chestnut oak to a galling cynipid. J. Chem. Ecol. 31, 151–166 (2005).

30. I. Kot, A. Jakubczyk, M. Karaś, U. Złotek, Biochemical responses induced in galls of three Cynipidae species in oak trees. Bull. Entomol. Res. 108, 494–500 (2018).

31. S. E. Hartley, The chemical composition of plant galls: are levels of nutrients and secondary compounds controlled by the gall-former? Oecologia 113, 492–501 (1998).

32. F. Ronquist, et al., Phylogeny, evolution and classification of gall wasps: the plot thickens. PLoS One 10, e0123301 (2015).

33. H. W. Kim, et al., NPClassifier: A Deep Neural Network-Based Structural Classification Tool for Natural Products. J. Nat. Prod. 84, 2795–2807 (2021).

34. I. Kot, C. Sempruch, K. Rubinowska, W. Michałek, Effect of (L.) galls on physiological and biochemical response of leaves. Bull. Entomol. Res. 110, 34–43 (2020).

35. I. Kot, K. Rubinowska, W. Michałek, CHANGES IN CHLOROPHYLL a FLUORESCENCE AND PIGMENTS COMPOSITION IN OAK LEAVES WITH GALLS OF TWO CYNIPID SPECIES (Hymenoptera, Cynipidae). Acta Sci. Pol. Hortorum Cultus 17, 147–157 (2018).

36. M. Calderón-Santiago, M. A. Fernández-Peralbo, F. Priego-Capote, M.D. Luque de Castro, MSCombine: a tool for merging untargeted metabolomic data from high-resolution mass spectrometry in the positive and negative ionization modes. Metabolomics 12, 1–12 (2016).

37. L. W. Sumner, et al., Proposed minimum reporting standards for chemical analysis Chemical Analysis Working Group (CAWG) Metabolomics Standards Initiative (MSI). Metabolomics 3, 211–221 (2007).

38. B. Zhao, Q. Liu, B. Wang, F. Yuan, Roles of Phytohormones and Their Signaling Pathways in Leaf Development and Stress Responses. J. Agric. Food Chem. 69, 3566–3584 (2021).

39. M. A. Fenn, J. J. Giovannoni, Phytohormones in fruit development and maturation. The Plant Journal 105, 446–458 (2021).

40. Z. Yuan, D. Zhang, Roles of jasmonate signalling in plant inflorescence and flower development. Curr. Opin. Plant Biol. 27, 44–51 (2015).

41. J. C. Schultz, P. P. Edger, M. J. A. Body, H. M. Appel, A galling insect activates plant reproductive programs during gall development. Sci. Rep. 9, 1833 (2019).

42. J. F. Tooker, A. M. Helms, Phytohormone Dynamics Associated with Gall Insects, and their Potential Role in the Evolution of the Gall-Inducing Habit. Journal of Chemical Ecology 40, 742– 753 (2014).

43. A. Raman, C. W. Schaefer, T. M. Withers, Biology, Ecology, and Evolution of Gall-inducing Arthropods (2005).

44. M. Kojima, et al., Highly sensitive and high-throughput analysis of plant hormones using MS-probe modification and liquid chromatography-tandem mass spectrometry: an application for hormone profiling in Oryza sativa. Plant Cell Physiol. 50, 1201–1214 (2009).

45. M. Tokuda, et al., Phytohormones related to host plant manipulation by a gall-inducing leafhopper. PLoS One 8, e62350 (2013).

46. L. Zhu, M.-S. Chen, X. Liu, Changes in Phytohormones and Fatty Acids in Wheat and Rice Seedlings in Response to Hessian Fly (Diptera: Cecidomyiidae) Infestation. Journal of Economic Entomology 104, 1384–1392 (2011).

47. C. Tayeh, B. Randoux, D. Vincent, N. Bourdon, P. Reignault, Exogenous Trehalose Induces Defenses in Wheat Before and During a Biotic Stress Caused by Powdery Mildew. Phytopathology® 104, 293–305 (2014).

48. , Advances in Insect Physiology (Elsevier, 2003).

49. Y.-N. Li, Y.-B. Liu, X.-Q. Xie, J.-N. Zhang, W.-L. Li, The Modulation of Trehalose Metabolism by 20-Hydroxyecdysone in Antheraea pernyi (Lepidoptera: Saturniidae) During its Diapause Termination and Post-Termination Period. J. Insect Sci. 20 (2020).

50. J. Weng, C. Chapple, The origin and evolution of lignin biosynthesis. New Phytologist 187, 273– 285 (2010).

51. M. Li, Y. Pu, A. J. Ragauskas, Current Understanding of the Correlation of Lignin Structure with Biomass Recalcitrance. Front Chem 4, 45 (2016).

52. J. Nakashima, F. Chen, L. Jackson, G. Shadle, R. A. Dixon, Multi-site genetic modification of monolignol biosynthesis in alfalfa (Medicago sativa): effects on lignin composition in specific cell types. New Phytol. 179, 738–750 (2008).

53. S. E. Brooks, J. D. Shorthouse, Developmental morphology of stem galls of Diplolepis nodulosa (Hymenoptera: Cynipidae) and those modified by the inquiline Periclistus pirata (Hymenoptera: Cynipidae) on Rosa blanda (Rosaceae). Canadian Journal of Botany 76, 365–381 (1998).

54. D. Wool, R. Aloni, O. Ben-Zvi, M. Wollberg, A galling aphid furnishes its home with a built-in pipeline to the host food supply. Proceedings of the 10th International Symposium on Insect-Plant Relationships, 183–186 (1999).

55. M. R. Samaan, A. A. Morsy, H. A. Kassem, Anatomical aspects of stem galls induced by Rhopalomyia spp. On their host plants. J. Environ. Sci. 45, 19–35 (2019).

56. Y. Zeng, M. E. Himmel, S.-Y. Ding, Visualizing chemical functionality in plant cell walls. Biotechnol. Biofuels 10, 263 (2017).

57. L. D. Salvador, T. Suganuma, K. Kitahara, H. Tanoue, M. Ichiki, Monosaccharide composition of sweetpotato fiber and cell wall polysaccharides from sweetpotato, cassava, and potato analyzed by the high-performance anion exchange chromatography with pulsed amperometric detection method. J. Agric. Food Chem. 48, 3448–3454 (2000).

58. J. S. Kim, G. Daniel, Immunolocalization of hemicelluloses in Arabidopsis thaliana stem. Part I: temporal and spatial distribution of xylans. Planta 236, 1275–1288 (2012).

59. J. B. Taft, D. R. Bissing, DEVELOPMENTAL ANATOMY OF THE HORNED OAK GALL INDUCED BY CALLIRHYTIS CORNIGERA ON QUERCUS PALUSTRIS (PIN OAK). American Journal of Botany 75, 26–36 (1988).

60. T. Tzfira, V. Citovsky, Agrobacterium: From Biology to Biotechnology (Springer, 2008).

61. C. I. Ullrich, R. Aloni, Vascularization is a general requirement for growth of plant and animal tumours. J. Exp. Bot. 51, 1951–1960 (2000).

62. R. Aloni, K. Pradel, C. Ullrich, The three-dimensional structure of vascular tissues in Agrobacterium tumefaciens-induced crown galls and in the host stems of Ricinus communis L. Planta 196 (1995).

63. A. Dolzblasz, A. Banasiak, D. Vereecke, Neovascularization during leafy gall formation on Arabidopsis thaliana upon Rhodococcus fascians infection. Planta 247, 215–228 (2018).

64. E. W. Nester, Agrobacterium: natureâ€^TM^s genetic engineer. Frontiers in Plant Science 5 (2015).

65. L. S. Thomashow, S. Reeves, M. F. Thomashow, Crown gall oncogenesis: evidence that a T-DNA gene from the Agrobacterium Ti plasmid pTiA6 encodes an enzyme that catalyzes synthesis of indoleacetic acid. Proc. Natl. Acad. Sci. U. S. A. 81, 5071–5075 (1984).

66. D. E. Akiyoshi, H. Klee, R. M. Amasino, E. W. Nester, M. P. Gordon, T-DNA of Agrobacterium tumefaciens encodes an enzyme of cytokinin biosynthesis. Proc. Natl. Acad. Sci. U. S. A. 81, 5994–5998 (1984).

67. D. E. Akiyoshi, et al., Cytokinin/auxin balance in crown gall tumors is regulated by specific loci in the T-DNA. Proc. Natl. Acad. Sci. U. S. A. 80, 407–411 (1983).

68. J. De Meutter, et al., Production of auxin and related compounds by the plant parasitic nematodes Heterodera schachtii and Meloidogyne incognita. Commun. Agric. Appl. Biol. Sci. 70, 51–60 (2005).

69. J. E. Jeon, et al., A Pathogen-Responsive Gene Cluster for Highly Modified Fatty Acids in Tomato. Cell 180, 176–187.e19 (2020).

70. M. Wang, et al., Sharing and community curation of mass spectrometry data with Global Natural Products Social Molecular Networking. Nat. Biotechnol. 34, 828–837 (2016).

71. P. Shannon, et al., Cytoscape: a software environment for integrated models of biomolecular interaction networks. Genome Res. 13, 2498–2504 (2003).

72. A. T. Aron, et al., Reproducible molecular networking of untargeted mass spectrometry data using GNPS. Nat. Protoc. 15, 1954–1991 (2020).

73. S. Suzuki, et al., High-throughput determination of thioglycolic acid lignin from rice. Plant Biotechnol. 26, 337–340 (2009).

74. J. Harholt, et al., ARABINAN DEFICIENT 1 is a putative arabinosyltransferase involved in biosynthesis of pectic arabinan in Arabidopsis. Plant Physiol. 140, 49–58 (2006).

75. A. Eudes, et al., Expression of a bacterial 3-dehydroshikimate dehydratase reduces lignin content and improves biomass saccharification efficiency. Plant Biotechnol. J. 13, 1241–1250 (2015).

76. L. Fang, et al., Loss of Inositol Phosphorylceramide Sphingolipid Mannosylation Induces Plant Immune Responses and Reduces Cellulose Content in Arabidopsis. Plant Cell 28, 2991–3004 (2016).

